# Divergent mitochondrial and metabolic adaptations shape selective vulnerability in ALS

**DOI:** 10.64898/2026.01.30.702552

**Authors:** Bianca A Cotto, Lizi Zhang, Benjamin Fait, Chris Peralta, Ece Kilic, Henrik Molina, Nathaniel Heintz, Eric F Schmidt

## Abstract

Neurodegenerative diseases like amyotrophic lateral sclerosis (ALS) exhibit striking cell-type selectivity, yet the basis for this vulnerability remains elusive. Here, we uncover that even closely related neurons can harbor distinct mitochondrial properties that shape their response to disease. Using TOM-Tag, a circuit-based AAV-based strategy for cell type–specific mitochondrial immunopurification from projection neurons, we performed integrative proteomic, metabolomic, transcriptomic, and functional analyses of mitochondria from ALS-vulnerable corticospinal projection neurons (CSPNs) and resilient corticothalamic projection neurons (CTPNs) in vivo. We discovered that CSPNs and CTPNs exhibit divergent mitochondrial profiles at baseline, despite sharing cortical layer and developmental origin. CTPNs were primed for antioxidant buffering and fatty acid metabolism, whereas CSPNs were enriched for oxidative phosphorylation components. In ALS, CTPNs employed mitochondrial flexibility and redox defense, whereas CSPNs exhibited respiratory failure and metabolic stress. These findings reveal that intrinsic mitochondrial programs vary even between similar neurons, and that this hidden layer of diversity may critically shape susceptibility to neurodegeneration. By enabling high-resolution access to mitochondria in defined neuronal circuits, TOM-Tag offers a powerful new lens for dissecting disease mechanisms and identifying cell-specific therapeutic targets.

## Introduction

The central nervous system (CNS) is composed of diverse populations of long-lived neurons with varying degrees of susceptibility to neurodegenerative diseases, although the basis for this phenomenon remains poorly understood. Amyotrophic lateral sclerosis (ALS), a disorder characterized by the selective loss of neurons in central motor pathways, epitomizes the cell type-specific nature of neurodegenerative diseases. In ALS, degeneration is observed in both layer 5b corticospinal projecting motor neurons (CSPNs) originating in the primary motor cortex (M1) and their downstream targets, lower motor neurons (LMN) located in the ventral horn of the spinal cord, which innervate muscle fibers. The loss of these cell populations impacts voluntary movement, leading to muscle weakness that progresses to debilitating and irreversible paralysis^1^ . The majority of ALS cases have no identifiable genetic cause, and for all cases, ALS remains incurable^2^ . Irrespective of the disease causation, the selective loss of motor neurons and preservation of neighboring neuronal cell populations is a hallmark feature of ALS pointing to the likelihood that intrinsic properties of motor neurons are paramount to their selective vulnerability.

A growing body of pathological and clinical evidence points to the importance of early cortical involvement in the onset and progression of ALS^3^ . In mouse strains generated to mimic ALS symptomology, loss of CSPNs has been reported and demonstrated to occur prior to LMN degeneration^4–9^. Together this emphasizes the need to investigate the molecular mechanism underlying CSPN vulnerability. Advances in cell type specific gene profiling techniques have shed light on the unique molecular signature of CSPNs. Our group and others have shown CSPNs are highly enriched for mitochondrial-related transcripts in contrast to whole brain and even other closely related cell populations^10–12^ . This suggests CSPNs may have a unique reliance on mitochondria. Interestingly, computational estimates of the bioenergetic requirements for MNs place them on the higher end of energy demands as they are extremely active, continuously firing to maintain posture or to sustain the firing patterns needed for muscle contraction during movements^13^. They also have quite large cell bodies with long axons, adding to their metabolic burden. Beyond bioenergetic pathways, we observed differences in the expression of genes involved in handling reactive oxygen species reflecting an additional inherent mitochondrial-related property of CSPNs^10^. Together these findings not only support the notion that corticospinal-projecting MNs have greater metabolic needs than other neuronal types but also suggest the likelihood of their bioenergetic constraints and mitochondrial properties contributing to their heightened vulnerability.

Mitochondria are multifaceted organelles which orchestrate various homeostatic functions in addition to supplying ATP, including modulation of Ca^2+^ flux and generation of essential building blocks for macromolecules such as lipids, proteins, DNA and RNA^14^. Importantly, mitochondria serve as sensors for various forms of cellular stress, including nutrient deprivation, oxidative stress, DNA damage, and endoplasmic reticulum stress. They coordinate the cellular adaptions behind whether a cell mounts a survival response or activates a cell death pathway. Neuronal fitness is reliant on the ability to maintain a steady supply of dynamic mitochondria able to respond to the intricate balance of bioenergetics, trafficking, quality control, fission/fusion, and biogenesis^15^ . In recent years the heterogeneity of mitochondria has become of increasing significance. Mitochondria derived from different tissue sources have been shown to have differing proteomes which implies tissue-specific mitochondrial dynamics^16–21^ . In the brain, distinctions have been made between the morphology, composition, and function of synaptic versus nonsynaptic mitochondria^22–25^ . Yet little is known about the extent of cell type-specific mitochondrial heterogeneity in the brain. Recent studies have uncovered distinct mitochondrial proteomes, lipid metabolism, Ca2+ buffering properties, and inter-organellar interactions between glia and major neuronal cell types^26^ . Understanding the extent to which mitochondria diversity exists in the brain could lead to important insights into brain health and neurodegeneration.

Mitochondrial pathology is a common observation in ALS and ALS model systems^27,28^ . Mitochondrial abnormalities and disrupted energy metabolism have been reported to occur even prior to onset of symptoms further underscoring the significance of mitochondrial integrity in ALS etiology^29–31^ . Why these changes in bioenergetics and mitochondria function seem to be occurring specifically in MNs is unknown. A better understanding of the intrinsics properties that govern CSPN’s mitochondrial dynamics and their response to diseases like ALS is needed.

Here, we sought to investigate the mitochondrial features of CSPNs and how they may compromise their survival in ALS. We implemented a multi-omics approach to assess the molecular, proteomic, and metabolomic features of two distinct L5b projection neuron subpopulations in M1 to identify unique cell specific molecular processes important for cellular vulnerability, resilience and mitochondrial fitness in ALS using the hSOD1-G93a mouse model^32^ . We developed and applied a strategy to facilitate the isolation of mitochondria by magnetic beads from whole tissue homogenates in a cell type-specific manner, termed TOM-Tag. By leveraging circuit-based neuronal targeting in mice with Cre recombinase expression restricted to L5b we could assess mitochondria from CSPNs and a neighboring, highly similar but ALS-resilient population of subcortically projecting neurons. This revealed the existence of divergent mitochondrial signatures between these closely related neuronal subpopulations and the impact these differences have in the cellular response to ALS.

## Results

### Projection-Specific Targeting of L5b Cortical Neurons for Translatome Profiling

CSPN and L5b corticothalamic projecting neurons (CTPNs) are closely related neighboring excitatory projection neurons with cell bodies found in layer 5b of the mouse cortex^33^ . They share classical molecular markers for long-distance projection neurons such as *Ctip2* and *Fezf2*. Yet they are molecularly distinct populations of cortical neurons located in discrete sublayers of L5b^10,33^ and are differentially vulnerable to ALS. Work with bacTRAP transgenic lines targeting cells in each population has shown that, contrary to CSPNs, the L5b CTPNs are resistant to degeneration in symptomatic hSOD^*^G93A mice, a widely used model of familial ALS^32^, and thus represent a valuable resilient population of cortical projection neurons^10^. To confirm this in non-bacTRAP mice, we took advantage of the selective expression of *Vat1l* in CTPNs and *Serpina3n* in CSPNs and measured the number of *Vat1l*-positive or *Serpina3n*-positive soma in the upper layer L5b of wild-type and hSOD1^*^G93A mice at 120 days of age (postnatal day 120, P120), a time point where limb paralysis is evident and cell loss is noted in cortex^4,7,9^. While we found a reduction in *Serpina3n*-expressing soma in M1, there was no change in the number of *Vat1l*-positive soma (**Figure 1a, b**). We reasoned that the comparison between the two L5b populations would distinguish genes and pathways that may predispose CSPNs to vulnerability and give rise to differential responses to ALS, and/or underlie selective neuronal degeneration.

**Figure 1.**
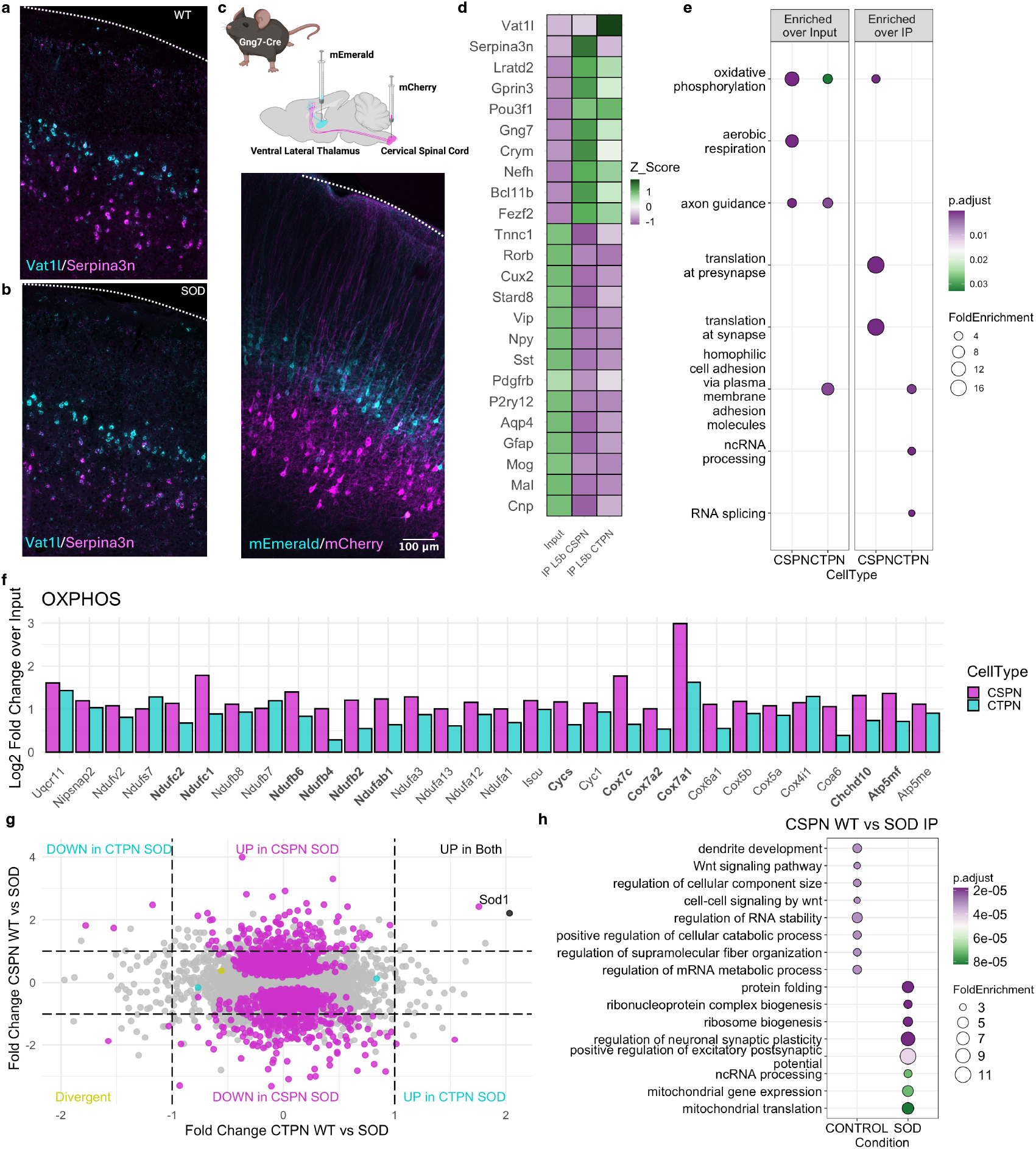
Cell type-specific translatome profiling of ALS-vulnerable and resilient L5b projection neurons. **a**,**b**, Representative confocal images of in situ hybridization showing *Vat1L*+ CTPN in upper L5b and *Serpina3n*+ CSPNs in lower L5b in M1 of WT (a) or SOD1G93A (b) mice at P120. **c**, Retrograde AAV-FLEX-mCherry injection into C4–C5 (CSPNs) or AAV-FLEX-mEmerald into VLT (CTPNs) in Gng7-Cre mice for circuit-based targeting of L5b populations in M1 shows mEmerald labeling restricted to cells in upper L5b (CTPNs) and mCherry expression in lower L5b (CSPNs). **d**, Heatmap displaying Z-scores for canonical glial and cortical neuron marker genes in CSPN and CTPN TRAP IPs relative to input. **e**, Top GO BP terms in CSPN- and CTPN-enriched genes relative to input (left) or relative to one another (right) **f**, Log_2_ fold change for CSPN-enriched genes contributing to the oxidative phosphorylation (OXPHOS) GO category in TRAP IPs relative to input based on differential expression analysis (padj < 0.05). Bold gene names show cell-specific expression in IPs. **g**, Scatter plot comparing SOD1G93A– mediated gene expression changes in CSPNs and CTPNs. Each point represents a gene with a mean counts-per-million (CPM) >100 across all samples. Log_2_ fold change (SOD1 vs. WT) in CTPNs is plotted on the x-axis; corresponding values for CSPNs are plotted on the y-axis. Genes significantly changed (padj<0.05) in CSPNs only (magenta), CTPNs only (cyan), or both cell types (black) are indicated. Dotted lines denote a log_2_ fold change threshold of ±1. **h**, Top GO terms enriched for genes significantly up- or downregulated in CSPNs in disease.

Previously, we used Gprin3-bacTRAP and Colgalt2-bacTRAP transgenic lines to access CSPNs or CTPNs, respectively^10^ . However, to obtain cleaner populations and reduce the number of transgenic lines needed, we took advantage of the distinct axonal projections of CSPNs and CTPNs and employed a retrograde viral approach to selectively target each population. Cre-dependent AAV vectors encoding either mCherry or mEmerald were injected bilaterally into either the cervical spinal cord (C4–C5), a distal target of CSPNs, or the ventrolateral thalamus (VLT), a distal target of layer 5b CTPNs, respectively, of Gng7-Cre driver mice, where Cre recombinase is restricted to all L5b neurons in cortex. Examination of M1 revealed mCherry expression restricted to lower L5b neurons and mEmerald expression in upper L5b neurons, consistent with the distribution of *Serpina3n* and *Vat1l* (**Figure 1c**) establishing this circuit-based targeting as viable approach for genetically targeting the CSPN and CTPN populations.

To investigate the unique properties of the L5b populations, we generated cell type-specific gene expression profiles for CSPN and CTPN using the viral Translating Ribosome Affinity Purification (vTRAP) approach in Gng7-Cre mice. The vTRAP approach has been demonstrated to capture cell type-specific translational profiles in any cell type engineered to express Cre recombinase in a spatiotemporally restricted fashion with high fidelity while allowing for more localized molecular profiling^34^. The vTRAP construct was packaged into virus coated with the retrograde AAV capsid^35^ to enable targeting projection populations by injection into the sites of axon terminations. Retro-AAV-FLEX-EGFPL10a, was injected bilaterally into C4-C5 or VLT. Anti-GFP immunostaining of M1 revealed bright cytoplasmic expression restricted to upper or lower L5b cells corresponding to CTPNs or CSPNs respectively (**Supplementary Figure 1a**).

For translational profiling, M1 was dissected from retro-AAV-FLEX-EGFPL10a injected mice and TRAP performed on tissue from individual animals to isolate tagged polysomes. Polysome-bound transcripts from CSPN and L5b CTPN neurons were analyzed by high-throughput RNA sequencing (RNA-seq). Samples from both L5b CTPN and CSPN IPs were enriched for classical projection neuron genes and depleted for glial markers relative to RNA isolated from whole tissue pre-IP M1 “input” samples, confirming cell type specificity of vTRAP isolations (**Figure 1d**). Unsupervised hierarchical clustering of all samples clearly separated the M1 input samples and the two neuronal populations into distinct clusters (**Supplementary Figure 1b**) which was confirmed by principal component analysis (PCA) of the top 500 variable genes globally distinguished the L5b CTPN IPs from the CSPN (**Supplementary Figure 1c**). Differential expression (DE) analysis identified 627 genes enriched (padj <0.05, log2FoldChange > 1 and baseMean > 100 log_10_CPM) in CSPNs over the input and 344 genes enriched in CTPNs over input (**Supplementary Figure 1d-e**). Gene ontology (GO) analysis for biological processes (BP) on the genes enriched in CSPNs confirmed the top pathways for this cell type are related to mitochondrial respiration (**Figure 1e** and **Supplementary Figure 1f**). Alternatively, these GO terms were enriched to a lesser extend in L5b CTPNs while GO terms related to adhesive molecules and neuron projection guidance were among their top significantly enriched in these cells (**Figure 1e** and **Supplementary Figure 1g**). We performed DE analysis to directly compare CSPN and L5b CTPN IPs and identified 1782 differentially expressed genes (DEG) (padj <0.05, log2FoldChange |0.5| and baseMean > 100 log_10_CPM) (**Supplementary Figure 1h**). GO analysis for BP in DEGs up in CSPNs over CTPNs also showed enrichment of mitochondrial-related processes like oxidative phosphorylation (OXPHOS) as well as pathways involved translation at synapses (**Figure 1e**). In contrast, DEGs in L5b CTPNs over CSPNs were related to ncRNA processing, RNA splicing, and ribonucleoprotein complex biogenesis (**Figure 1e**). Among the genes driving this enrichment for OXPHOS GO term in CSPN were complex I and complex IV subunits as well as *Chchd10*, a previously identified ALS-related gene^36,37^ (**Figure 1f**).

### Translatome Profiling of ALS-Vulnerable and -Resilient Layer 5b Projection Neurons

To investigate molecular mechanisms of ALS-related degeneration in vulnerable CSPNs, we used vTRAP to compare cell type-specific expression changes between CSPNs and the resilient L5b CTPNs. Gng7-Cre mice were crossed to hSOD1^*^G93A mice and at P35 retro-AAV-FLEX-EGFPL10a was injected into the C4-C5 spinal cord or into the VLT of Cre+ animals expressing the mutant SOD transgene (SOD) as well as non-SOD littermates (WT). M1 was harvested for TRAP at the symptomatic age of P120 when motor deficits were observed in rotarod performance^10,38,39^ (**Supplementary Figure 1i**). While PCA of the top 500 variable genes globally distinguished the L5b CTPN IPs from the CSPN regardless of the mutant SOD transgene expression wildtype and SOD IPs showed a sub-group separation in CSPNs but no sub-groups were visible within the L5b CTPN samples (**Supplementary Figure 1j**). Mapping TRAP-seq reads to the human SOD gene showed the mutant SOD transgene was expressed at similar levels in each cell type (**Supplementary Figure 1k**).

Next, we identified genes that showed a significant change in expression in disease by performing DE analysis between WT and SOD samples. Vulnerable CSPNs showed a robust response to SOD1-G93A expression, with 1264 genes differentially regulated (571 up-regulated and 693 down-regulated; **Figure 1g**). In contrast, only four genes were significantly altered in disease in the ALS-resilient L5b CTPN, (two up-regulated and two downregulated; **Figure 1g**). Only *Sod1* was significantly up-regulated in both cell types and there were no commonly down-regulated genes (**Figure 1g**). GO analyses on DE genes from CSPN cells revealed genes related to protein folding, ribonucleoprotein complex biogenesis, mitochondrial gene expression and mitochondrial translation were up-regulated, while genes associated with dendrite development, Wnt signaling, and RNA stability were down-regulated (**Figure 1h**). These results indicated a relatively robust molecular response to SOD1^*^G93A in vulnerable CSPNs, but not L5 CTPNs, that pointed to a triggering of mitochondrial gene expression machinery in response to hSOD1^*^G93A expression.

### TOM-Tag Enables Cell Type-Specific Mitochondrial Isolation from L5b Projection Neurons

To determine the significance of enrichment of mitochondrial-related transcripts in CSPNs, we sought to isolate, characterize, and compare mitochondria from CSPNs and L5b CTPNs. To develop a viral vector that would enable the isolation of intact mitochondria from specific neuronal cell types, we subcloned the cDNA of mEmerald-tagged outer mitochondrial membrane protein TOMM20 (mEmerald-TOMM20) into the Cre-dependent pAAV-EF1a-DIO-EGFPL10a plasmid, replacing EGFPL10a. The resulting virus had a double-floxed inverse open reading frame to allow for the conditional expression of the mEmerald-TOMM20 fusion protein in the presence of Cre recombinase. This vector is hereafter referred to as pAAV-TOM-Tag (**Figure 2a**). Once incorporated into the outer mitochondrial membrane, the TOMM20-mEmerald protein allows for rapid immunopurification of intact mitochondria from whole tissue homogenates using anti-eGFP-coated magnetic beads (**Figure 2a**). To confirm Cre-dependent expression of the mEmerald-TOMM20 construct, we co-transfected HEK293T cells with pAAV-TOM-Tag and pAAV-Ef1a-mCherry-IRES-Cre. Immunofluorescent imaging of transfected cells and western blotting of cell lysates revealed that expression of the mEmerald-TOMM20 fusion protein was restricted only to cells expressing mCherry and thus contained the Cre plasmid (**Supplementary Figure 2a, b**). In these cells, the mEmerald-TOMM20 fusion protein showed the expected molecular weight (**Supplementary Figure 2c**). MitoTracker staining of co-transfected cells showed colocalization of mEmerald-TOMM20 fusion protein with MitoTracker consistent with incorporation of the transgene into intact mitochondria (**Supplementary Figure 1d**).

**Figure 2.**
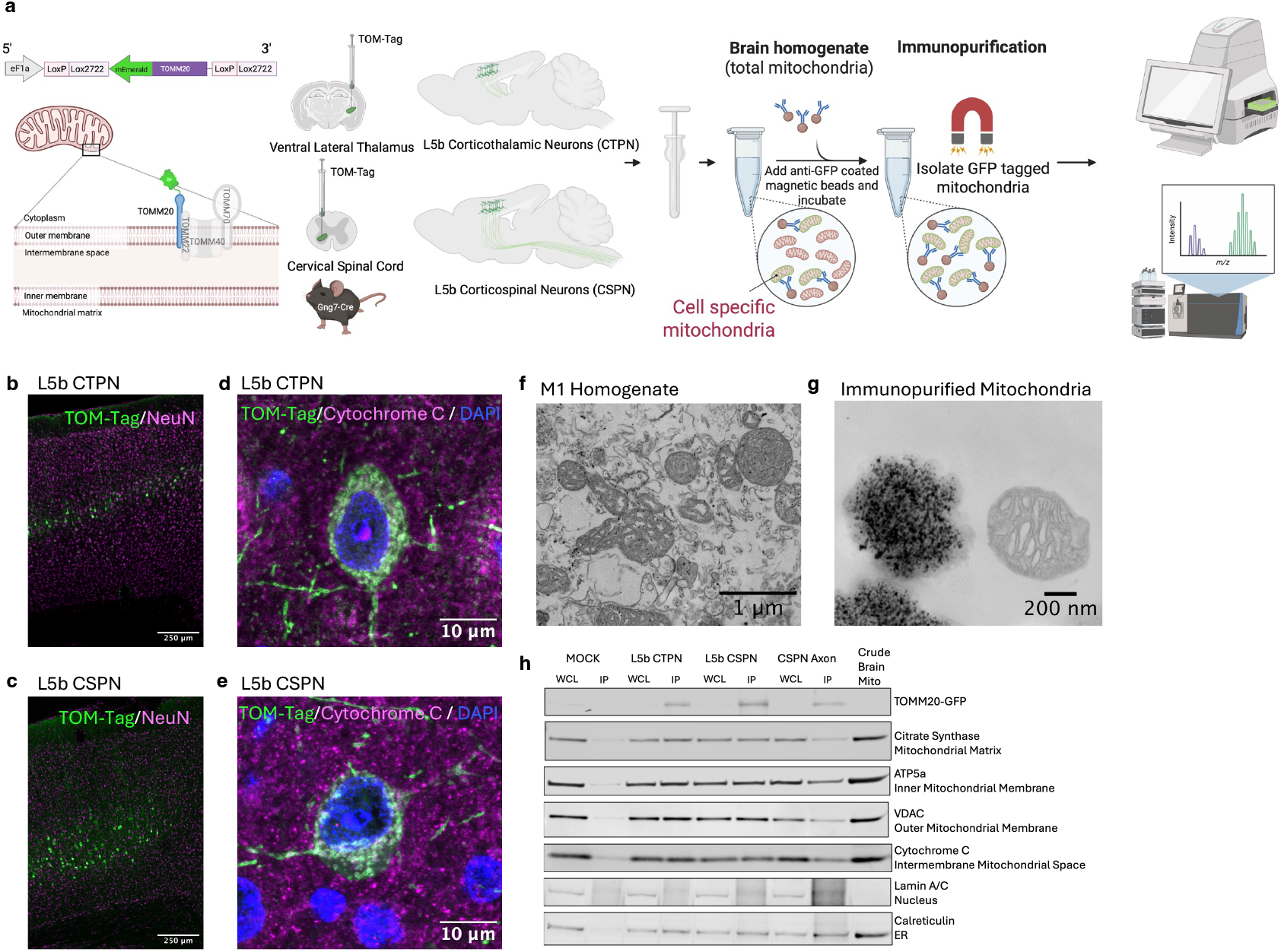
TOM-Tag enables cell-type-specific mitochondrial isolation from L5b projection neurons. **a**, Schematic overview of the TOM-Tag approach for cell type-specific mitochondrial profiling. The TOM-Tag construct encodes a Cre-dependent mEmerald-TOMM20 fusion protein expressed under the EF1α promoter. Upon retrograde AAV delivery into distal projection targets of CSPNs or CTPNs in Gng7-Cre mice, the fusion protein localizes to the outer mitochondrial membrane in Cre-expressing neurons. Mitochondria are subsequently isolated from M1 tissue homogenates via immunoprecipitation using anti-GFP–conjugated magnetic beads, enabling downstream proteomic and metabolic analyses. **b–c**, Immunofluorescence images of M1 showing laminar distribution of TOM-Tag (GFP, green) expression following retrograde AAV injection into VLT (CTPN, **b**) or cervical SC (CSPN, **c**) of Gng7-Cre mice. TOM-Tag signal localizes to distinct sublayers of deep layer 5 corresponding to each projection neuron population. Neuronal nuclei are stained with NeuN (magenta). **d–e**, High magnification confocal images of TOM-Tag (green) and Cytochrome C (magenta) immunolabeling in M1 confirm mitochondrial membrane localization of the fusion protein in CSPNs and CTPNs. Sections were counterstained with DAPI (blue). **f–g**, Transmission electron micrographs showing mitochondria in pre-IP whole tissue homogenate of M1 (f) and anti-GFP–conjugated magnetic beads bound to TOM-Tag– labeled mitochondria following immunocapture (g) in Gng7-Cre TOM-Tag-injected mice. **h**, Western blots for TOM-Tag fusion protein (mEmerald-TOMM20), mitochondrial markers (VDAC, Cytochrome C, ATP5a, Citrate Synthase), nuclear marker (Lamin A/C), and ER marker (Calreticulin) from whole-cell lysates (WCL) and immunoprecipitated (IP) fractions for CSPNs CTPNs from TOM-Tag injected Gng7-Cre mice, and mock controls (no TOM-Tag virus), axonal CSPN mitochondria isolated from midbrain tissue, and crude cortical mitochondria obtained by differential centrifugation.

To confirm the feasibility of the AAV-TOM-Tag approach for the isolation of mitochondria from genetically defined neuronal population within the mouse cortex, pAAV-TOM-Tag was packaged into the retrograde AAV vector (retro-AAV-TOM-Tag). Following bilateral injection of retro-AAV-TOM-Tag was into the cervical spinal cord or VLT of Gng7-Cre mice, anti-EGFP immunostaining demonstrated labeled cells in sublayers of M1 corresponding to CSPNs or CTPNs, respectively (**Figure 2b, c**). Double-labeling with anti-GFP and anti-Cytochrome C confirmed mitochondrial localization of the TOM-Tag transgene in both cell types (**Figure 2d, e**). Labeled mitochondria were isolated by immunopurification from dissected M1 homogenates using anti-GFP coated magnetic beads. Electron microscopy (EM) visualization of immunoprecipitates (IP) confirmed binding of the anti-GFP-coated beads to the outer membrane of intact mitochondria (**Figure 2f, g**). Western blot analysis of whole M1 homogenates and IPs from retrograde-targeted CSPN and CTPN samples showed specific enrichment for the fusion protein mEmerald-TOMM20 and the mitochondrial markers VDAC, Cytochrome C, ATP5a and Citrate Synthesis compared to a mock IP performed on mice which did not receive the TOM-Tag injection. The nuclear marker Lamin A/C was depleted in the IPs while the endoplasmic reticulum (ER) marker, calreticulin, was detected in the IP fraction, which is consistent with the fact that mitochondria can form contacts with the ER in neurons^40,41^ (**Figure 2h**). Together, these results show that TOM-Tag allows for the isolation of intact mitochondria from mouse brain.

## CSPN and CTPN Mitochondria Exhibit Divergent Bioenergetic Function and Metabolic Profiles

To test whether TOM-Tag isolated mitochondria were functionally competent, we performed a Seahorse Mito Stress Test using succinate plus rotenone to drive electron flow through Complex II. Sequential additions of oligomycin, FCCP, and antimycin A allowed assessment of key respiratory parameters. Isolated mitochondria from both CSPN and L5b CTPN responded robustly to these modulators, confirming preserved respiratory control. In WT Gng7-Cre mice, basal oxygen consumption rate (OCR) was similar in CSPN and CTPN mitochondria (**Figure 3a**), while maximal OCRs were higher in CTPN (**Figure 3b**). In SOD1^*^G93A mice, mitochondria from CSPNs displayed significantly lower basal and maximal OCR relative to WT CSPN and to CTPN SOD, whereas CTPN SOD increased respiration, with FCCP-stimulated OCR reaching similar maximum as their WT counterparts (**Figure 3a, b**). Because assays were performed with succinate plus rotenone, these differences reflect Complex II and downstream electron transport capacity rather than Complex I–dependent flux.

**Figure 3.**
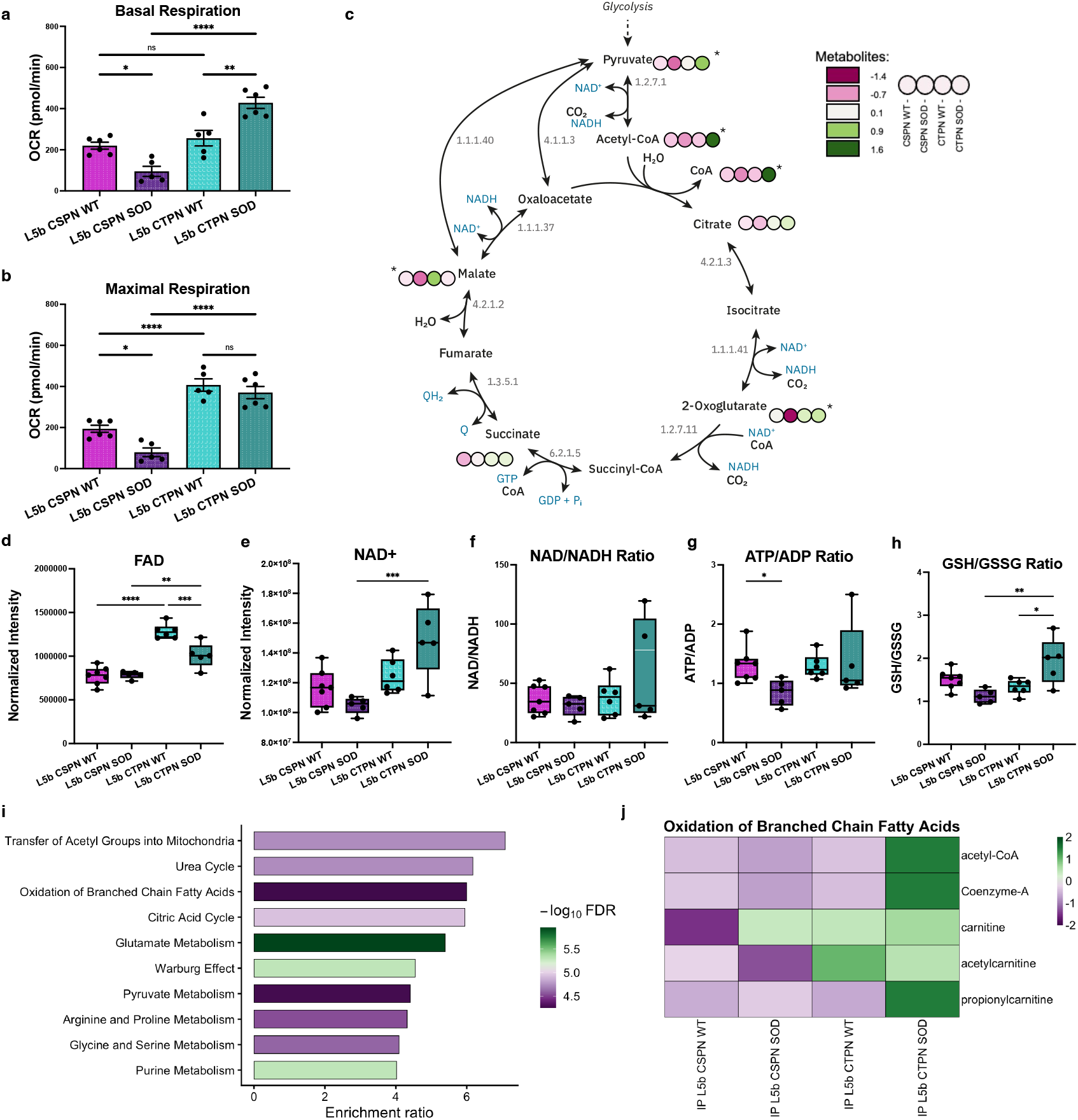
Functional divergence of CSPN and CTPN mitochondria revealed by respiration and metabolomics. **a–b**, Bar graphs showing quantification (mean ± s.e.m) of mitochondrial respiration in CSPNs and CTPNs from WT and SOD1G93A mice using the Seahorse Mito Stress Test where isolated mitochondria were fueled with succinate and rotenone to assess Complex II–dependent oxygen consumption. Basal OCR measured prior to drug injection (a) and maximal OCR measured following FCCP addition (b) are plotted. Each point represents a biological replicate. Statistical significance was assessed using one-way ANOVA with multiple comparisons. ^*^p = 0.01, ^**^p = 0.001, ^***^p = 0.0001, ^****^p = 0.00001, n.s = not significant. **c**, Diagram of the tricarboxylic acid (TCA) cycle overlaid with z-scored metabolite abundance values from LC–MS profiling of immunoprecipitated mitochondria from CSPNs and CTPNs in WT and SOD1G93A mice. Each circle represents a metabolite detected in the TCA pathway; color intensity reflects the normalized abundance (z-score) for each experimental group. Generated using MetaboMAPS^42^ . **d–e**, Box plots of normalized metabolite abundance measured by LC–MS in mitochondria immunoprecipitated from CSPNs and CTPNs in WT and SOD1^*^G93A mice for flavin adenine dinucleotide (FAD, d) and NAD^+^ (e). **f–h**, Box plots pf metabolic ratios derived from LC–MS quantification of metabolites in immunoprecipitated mitochondria from CSPNs and CTPNs from WT and SOD1^*^G93A mice showing NAD^+^/NADH ratio calculated from normalized abundance values, used as an indicator of mitochondrial redox state (f), ATP/ADP ratio calculated from normalized abundance values, representing phosphorylation potential and energetic state (g), and GSH/GSSG ratio derived from reduced and oxidized glutathione abundances, reflecting mitochondrial redox buffering capacity (h). **i**, Pathway enrichment analysis of metabolites that differed significantly across experimental groups, based on one-way ANOVA (p < 0.05). Metabolites were annotated to KEGG pathways, and enrichment analysis was performed to identify overrepresented metabolic processes among the group-wise variable metabolites. **j**, Heatmap showing z-scored abundance of metabolites annotated to the KEGG pathway “oxidation of branched-chain fatty acids,” measured by LC–MS in immunoprecipitated mitochondria from CSPNs and CTPNs across ALS genotypes. Z-scores were calculated across all samples for each metabolite.

To contextualize these functional differences, we quantified 111 polar metabolites in IPed mitochondrial preparations from CSPNs and L5b CTPNs in Gng7-Cre::SOD1G93A mice and WT littermate controls by ZIC-pHILIC LC–MS. One-way ANOVA identified 68 metabolites that differed across the four groups (FDR q<0.05). Alpha-ketoglutarate (α-KG) was selectively reduced in CSPN SOD mitochondria compared to CSPN WT and CSTN SOD groups, suggesting constrained TCA throughput (**Figure 3c**). In contrast, CTPN SOD samples exhibited higher levels of several TCA-linked intermediates, including pyruvate, acetyl-CoA, and CoA, compared to CTPN WT and CSPN SOD (**Figure 3c**). CSPNs also showed a diminshed α-KG /glutamate ratio in SOD, while both aspartate/glutamate and aspartate/malate increased, indicating a potential shift at the α-KG/OAA step toward glutamate and aspartate, leaving α-KG and malate relatively scarce (**Supplementary Figure 3a-c**). CSPN mitochondria did not show any change in FAD in disease, however FAD was higher in CTPN than CSPN overall, with a disease-related decrease within CTPN that still remained above CSPN levels (**Figure 3d**). Mitochondrial NAD^+^ levels were elevated in CTPN SOD compared to CTPN WT, although this increase did not reach statistical significance. However, NAD^+^ was significantly higher in CTPN SOD relative to CSPN SOD (**Figure 3e**). NADH levels and the NAD^+^/NADH ratio remained unchanged, indicating greater availability of oxidized NAD^+^ in CTPNs under disease conditions while maintaining redox balance (**Figure 3f**). Importantly, the ATP/ADP ratio was reduced in CSPN SOD relative to all other groups, indicating diminished phosphorylation potential (**Figure 3g**). Furthermore, redox buffering diverged by cell type in that GSH/GSSG increased in CTPN SOD but decreased in CSPN SOD (**Figure 3h**), consistent with enhanced antioxidant capacity in resilient CTPN and oxidative stress in vulnerable CSPN.

Using the ANOVA-significant metabolite list for over-representation analysis across all samples, we observed enrichment for fatty-acid oxidation, pyruvate metabolism, and glutamate metabolism (**Figure 3i**). Examination of metabolites annotated to “Oxidation of Branched-Chain Fatty Acid” revealed generally lower carnitine species (carnitine and acetylcarnitine) in CSPN relative to CTPN in WT, although carnitine was increased to comparable levels in the presence of SOD (**Figure 3j**). However, this was accompanied by a decrease in acetylcarnitine in CSPN SOD samples, while CSTNs showed increased acetyl-Co-A, Coenzyme A, and propionylcarnitine in SOD compared to WT. Propionylcarnitine elevations in CTPN SOD suggest increased odd-chain or branched-chain acyl flux toward propionyl-CoA and succinyl-CoA entry, consistent with greater fatty acid utilization. CTPN SOD showed higher FAD along with carnitine species changes, a pattern consistent with increased β-oxidation capacity that was not evident in the vulnerable CSPNs. These data taken together with the respiration profiles highlight fundamental differences in mitochondrial function between cell types where CTPNs appeared to enhance energy-producing pathways and redox buffering in disease, whereas the vulnerable CSPNs had reduced respiratory capacity, lower ATP/ADP, impaired α-KG availability, and weaker antioxidant status.

### Proteomic Landscape of Mitochondria from L5b Projection Neurons

Next, we determined whether functional differences between CSPN and CTPN mitochondria were reflected in their proteomic composition. To do this, we performed label-free data-independent acquisition (DIA)-based mass spectrometry on IP mitochondria from each cell type. As a reference, we included crude mitochondrial preparations isolated from M1 cortex by differential centrifugation, representing the bulk mitochondrial proteome of this brain region. Across all samples, we detected a total of 4,776 proteins. The number of proteins identified varied by sample origin, with 2,148 proteins detected across all conditions with 520 of these classified as mitochondrial according to MitoCarta3.0 (**Figure 4a**). Notably, 62 mitochondrial proteins were uniquely detected in the crude mitochondrial fraction, 27 in CTPN IP samples, and 6 in CSPN IP samples. CSPN IP samples yielded the fewest total proteins. To assess mitochondrial specificity and validate enrichment over background, we compared CSPN IP and CTPN IP samples to mock IP controls and identified 970 proteins differentially expressed in CSPN IPs and 880 in CTPN IPs (**Supplementary Figure 4a, c**). GO analysis of the cellular component category revealed significant enrichment for mitochondrial-associated terms including mitochondrial protein-containing complex, respirasome, and mitochondrial matrix in the CSPN and CTPN IP samples, but not in mock IP controls, confirming the specificity of the mitochondrial pulldown approach (**Supplementary Figure 4b, d**).

**Figure 4.**
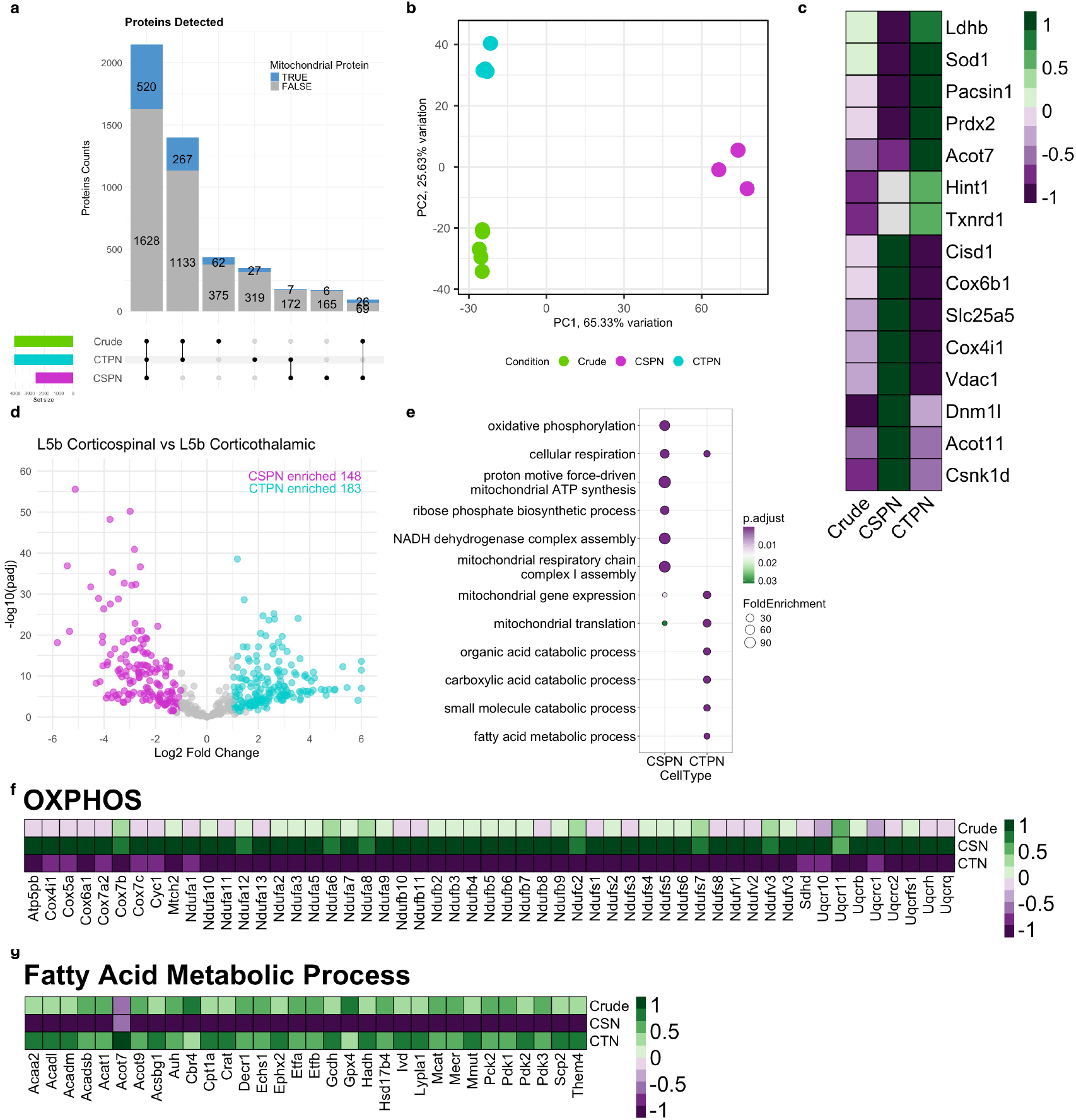
Proteomic profiling of CSPN and CTPN mitochondria reveals distinct metabolic programs. **a**, UpSet plot showing the intersection of proteins detected in CSPNTOM-Tag IP, CTPN TOM-Tag IP, and crude cortical mitochondrial preparations. Vertical bars indicate the number of proteins shared or unique to each sample type, with connected dots below denoting the data set membership. Proteins annotated as mitochondrial (based on MitoCarta3.0) are shown in light blue while non-mitochondrial proteins are shown in grey. **b**, PCA plot based on normalized abundance of all proteins shared across CSPN IP, CTPN IP, and crude cortical mitochondrial samples. **c**, Heatmap showing proteins most enriched in CSPNs (center) or CTPNs (right) relative to crude cortical mitochondria (left). Values represent Z-score of protein abundance for each sample. **d**, Volcano plot comparing mitochondrial protein abundance between CSPN IP and CTPN IP samples, limited to proteins annotated in MitoCarta3.0. Proteins significantly enriched (q < 0.05, |log_2_FC| ≥ 1) in CSPN mitochondria are shown in magenta while those enriched in CTPN mitochondria are shown in cyan. Non-significant proteins are shown in grey. The total number of significantly differentially expressed proteins in each direction is displayed in the upper right corner of the plot. **e**, GO enrichment analysis for biological process of MitoCarta3.0-annotated proteins differentially expressed between CSPN and CTPN IPs from d. **f**, Heatmap of z-scores of abundance values for proteins within the OXPHOS GO term, across crude cortical mitochondria, CSPN IPs, and CTPN IPs. **g**, Heatmap of z-scored abundance values for proteins associated with fatty acid metabolic processes across all groups.

Principal component analysis distinguished CSPN-IP and CTPN-IP proteomes from both crude mitochondrial preparations and each other, suggesting that each L5b projection neuron subtype harbored a distinct mitochondrial proteomic profile (**Figure 4b**). To identify proteins enriched in each neuronal subtype relative to total cortical mitochondria, we performed differential expression analysis between each IP sample (CSPN or CTPN) and the crude fraction. For CSPNs, the top enriched proteins included electron transport chain complex IV subunits (COX4I1 and COX6B1) and components of the permeability transition pore complex (VDAC1 and SCL25A5) (**Figure 4c**). CTPN-enriched proteins included antioxidant enzymes (SOD1, PRDX2 and TXNRD1) and ACOT7, an acyl-CoA hydrolase involved in fatty acid metabolism (**Figure 4c**).

To directly compare mitochondrial composition between the two projection neuron types, we performed differential analysis between CSPN and CTPN IP samples. Focusing on the subset of mitochondrial proteins (as defined by MitoCarta3.0) that showed an absolute fold change ≥1 and q-value <0.05, we identified 158 proteins enriched in CSPN and 183 in CTPN (**Figure 4d**). GO enrichment analysis of DE proteins revealed that oxidative phosphorylation pathways were significantly enriched in CSPNs, as was seen in the gene expression data, whereas fatty acid metabolic processes were enriched in CTPNs, consistent with a baseline preference for fatty-acid supported energetics in the metabolomics data (**Figure 4e-g**).

### Multi-Omics Comparison Identifies Cox7a1 as a Mitochondrial Protein Unique to CSPNs

To assess the concordance between transcriptional and mitochondrial proteomic profiles in L5b projection neurons, we compared our proteomic dataset with OXPHOS-related genes that were significantly enriched in CSPNs over CTPNs in the TRAP data sets (**Figure 1f** and **Figure 5a**). Overall, there was strong agreement between transcript and protein relative abundance levels, suggesting that key aspects of mitochondrial identity are transcriptionally encoded in both CSPNs and CTPNs. A small number of proteins, including NIPSNAP2, CYCS, and COA6, deviated from this trend, displaying discordance between mRNA expression and protein abundance, potentially reflecting cell type differences in translational regulation, protein turnover, or mitochondrial import. Of particular interest, COX7A1, which was significantly enriched in CSPNs at the transcript level was uniquely detected in the CSPN mitochondrial IP at the protein level (**Figure 5a**). This protein was not observed in either the CTPN IP samples or the crude cortical mitochondrial fraction, underscoring its specificity in CSPN mitochondria. To independently validate this finding, we performed RNAscope® in situ hybridization and confirmed the *Cox7a1* transcript co-expressed within *Serpina3n*+ CSPNs but not *Vat1l* CSTNs in the mouse M1 (**Figure 5b**).

**Figure 5.**
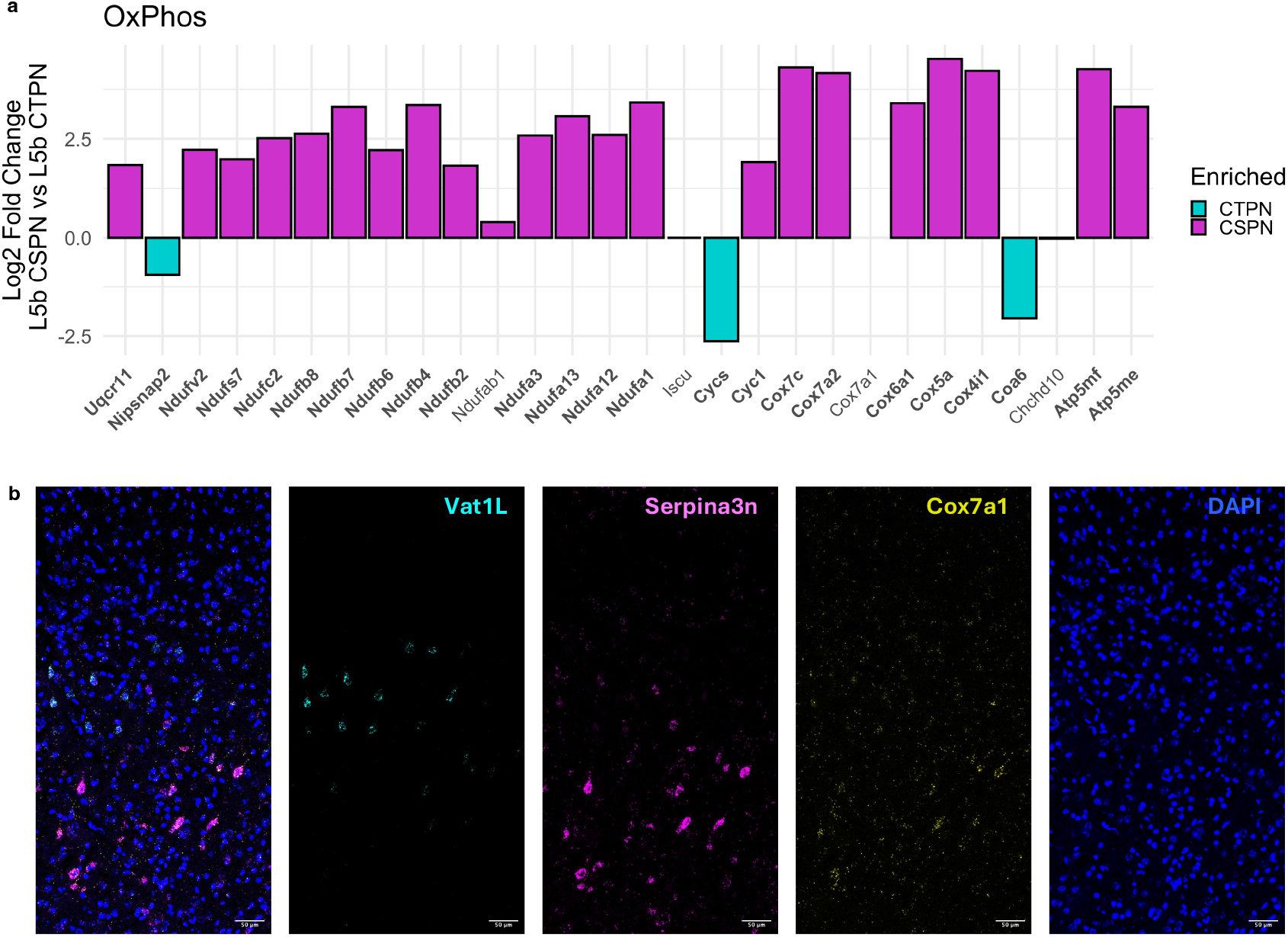
Cox7a1 is a CSPN-specific mitochondrial protein. **a**, Bar plot showing log_2_ fold change in protein abundance (CSPN vs. CTPN) for genes associated with OXPHOS that were significantly enriched at the transcript level in CSPN TRAP IPs (from Figure 1f). Protein names shown in bold were also significantly differentially expressed at the protein level. Notably, COX7A1 was not detected in CTPN TOM-Tag IPs. **b**, Representative confocal images from in situ hybridization in M1, showing labeling for *Vat1l* (cyan), *Serpina3n* (magenta), *Cox7a1* (yellow), and DAPI (blue). Left panel is an overlay of all probes.

### Cell-Type Specific Mitochondrial Responses to ALS Pathology

To determine how ALS-associated pathology alters mitochondrial composition in vulnerable and resilient neurons, DIA-based proteomic profiling was performed on TOM-Tag isolated mitochondria from CSPNs and CTPNs from SOD1^*^G93A Gng7-Cre mice and WT littermates at P120. We examined disease-related changes in crude M1 cortical mitochondrial preparations as a bulk tissue reference. Principal component analysis revealed clear separation of CSPNs and CTPNs along PC1, while PC2 further distinguished CSPN samples by genotype, similar to the TRAP data (**Supplementary Figure 1k**), once again indicating a robust disease-associated signature in CSPN mitochondria (**Figure 6a**). In contrast, CTPN samples showed no genotype-based separation, suggesting a more modest proteomic shift in response to ALS pathology or a heterogeneous response within this cell type (**Figure 6a**).

**Figure 6.**
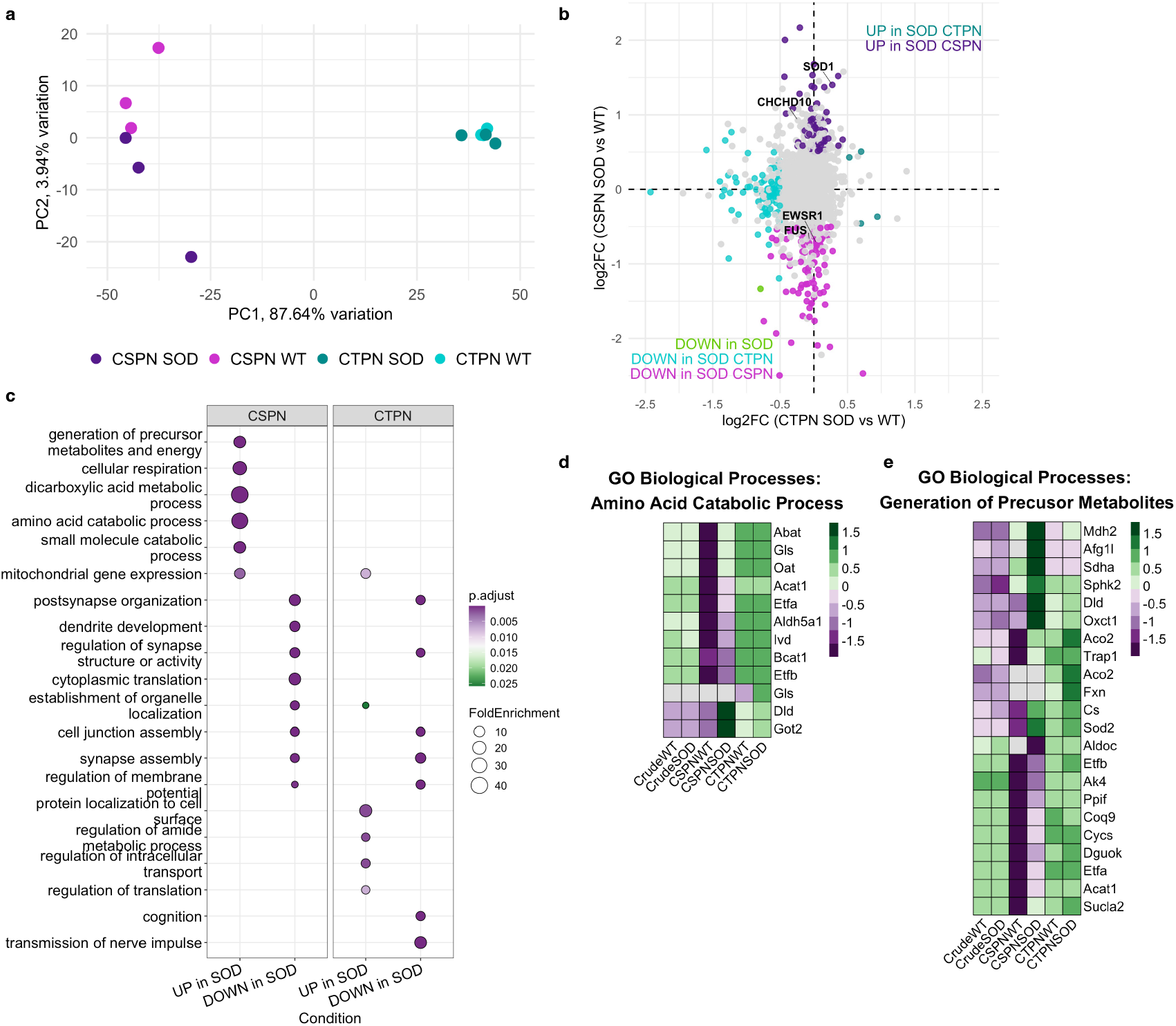
ALS-associated mitochondrial remodeling is cell-type specific. **a**, PCA plot of mitochondrial proteomic profiles from CSPN and CTPN TOM-Tag IP samples from M1 of WT and SOD1^*^G93A Gng7-Cre mice. Samples separated by cell type along PC1 (x-axis), with additional disease genotype-driven separation observed in CSPNs along PC2 (y-axis). **b**, Scatter plot comparing log_2_ fold changes in mitochondrial protein abundance (SOD vs. WT) in CTPNs (x-axis) and CSPNs (y-axis). Proteins significantly upregulated in both cell types (black), downregulated in both cell types (green), significantly upregulated only in CSPNs (dark purple) or CTPNs (dark cyan), or significantly downregulated only in CSPNs (magenta) or CTPNs (cyan) are indicated. **c**, GO biological process enrichment analysis of DE proteins in CSPN and CTPN IPs under disease conditions. **d–e**, Heatmaps showing z-scores of abundances of mitochondrial proteins annotated to the GO terms “amino acid catabolic process” (**d**) and “generation of precursor metabolites and energy” (**e**) across crude, CSPN, and CTPN mitochondrial samples from WT and SOD1^*^G93A mice.

Differential expression analysis of crude M1 mitochondria revealed no statistically significant changes between WT and SOD samples, although SOD1 itself approached significance (**Supplementary Figure 5a**). In contrast, CSPN mitochondria displayed a robust disease-associated signature, with 70 proteins significantly upregulated and 94 downregulated in SOD compared to WT (**Figure 6b** and **Supplementary Figure 5b**). In CTPNs, only 15 proteins were upregulated however 140 were downregulated in SOD1^*^G93A animals (**Figure 6b** and **Supplementary Figure 5c**). Notably, SOD1 was significantly increased in CSPN mitochondria but not in CTPNs (**Figure 6b** and **Supplementary Figure 5b, c**).

GO analysis of the differentially expressed mitochondrial proteins revealed distinct functional changes in each population (**Figure 6c**). In CSPNs, SOD-upregulated proteins were enriched for terms related to cellular respiration, amino acid metabolism, and dicarboxylic acid metabolic processes, while downregulated proteins were enriched for synaptic pathways, including postsynapse organization and dendrite development. In CTPNs, SOD-upregulated proteins were enriched for organelle organization and regulation of amide metabolic processes, whereas synaptic terms were again among the downregulated pathways, reflecting a shared vulnerability in synaptic function. To better understand the disease-associated metabolic response in CSPNs, we focused on the GO terms “amino acid catabolic process” and “generation of precursor metabolites and energy,” which were among the most enriched categories in CSPNs under SOD conditions (**Figure 6d, e**). Interestingly, these same pathways were more broadly enriched in CTPNs relative to CSPNs under baseline conditions, suggesting that CTPNs may be intrinsically better equipped to use alternate energy sources like fatty acids. In contrast, CSPNs appear to activate these programs only in response to disease stress. Several proteins within these pathways, including GOT2, MDH2, SLC25A10, GLUD1, IDH3, and the OGDH complex (OGDHL, OGDH, DLST, DLD), were significantly upregulated in CSPNs in the SOD condition relative to both WT CSPNs and CTPNs. This is consistent with upregulated malate– aspartate shuttle activity and in line with the metabolite ratios shown above revealing lower α-ketoglutarate to glutamate and higher aspartate to glutamate and aspartate to malate (**Supplementary Figure 3b-d**). Additionally, these proteomic shifts mirror our metabolomics findings, which suggest enhanced fatty acid utilization in both cell types during disease, but to a greater extent in CTPNs. Together, these data suggest that CTPNs are primed for substrate switching, whereas CSPNs depend on stress-induced compensatory remodeling, potentially limiting the effectiveness of their adaptation.

## Discussion

Understanding why certain neuronal populations degenerate in ALS while others remain resilient is a longstanding challenge in neurodegeneration research. In this study, we leveraged the closely related but differentially vulnerable corticospinal (CSPN) and corticothalamic (CTPN) projection neurons of layer 5b to investigate the molecular and mitochondrial features that influence selective vulnerability. Despite sharing classical molecular markers and occupying adjacent cortical sublayers, CSPNs are consistently among the earliest and most severely affected neuronal populations in ALS^4–7^, whereas CTPNs are spared into late stages of disease despite the presence of pathogenic mutant SOD1 expression. By exploiting their divergent projection targets and using a combination of vTRAP-based translatome profiling, TOM-Tag mitochondrial proteomics, and targeted metabolomics, we define, for the first time, the cell type-specific mitochondrial landscapes of these neurons and uncover fundamental differences in baseline state, disease response, and metabolic flexibility.

Our results uncover a distinct mitochondrial profile underlying CSPN vulnerability. Despite their molecular enrichment for oxidative phosphorylation components, CSPNs do not outperform resilient neurons like CTPNs under basal conditions; instead, they appear to operate near their metabolic ceiling. This disconnect may stem from several factors, including post translational regulation of OXPHOS activity, differences in substrate availability or preference, or the specific conditions of our Seahorse assay, which relied on Complex II driven respiration. Regardless of the mechanism, this rigid configuration leaves CSPNs poorly equipped to manage the redox imbalance and energetic demands imposed by disease. In ALS, this fragility manifests as impaired respiratory capacity and compromised antioxidant buffering, suggesting that CSPNs are unable to effectively rewire their mitochondrial machinery when challenged. These findings shift the perspective from viewing vulnerable neurons as simply damaged in disease to recognizing them as metabolically constrained even in health. This concept may help explain their unique susceptibility in ALS and inform future cell specific therapeutic strategies.

Our development of TOM-Tag enabled the highly specific isolation of mitochondria from defined projection neuron populations in vivo, facilitating the first multi-omics analyses of mitochondria from ALS-vulnerable and resilient cortical neurons. Previous cell type-specific mitochondrial isolation strategies, such as MITO-Tag^40^, MitoTag ^26,43,44^, and AAV-based outer membrane HA-tagged^45^ (e.g., AAV-mitoTag), have been instrumental in elucidating mitochondrial diversity and provided important insights into mitochondrial biology within genetically defined populations. Building on these foundational tools, we adapted a Cre-dependent mitochondrial labeling strategy to achieve retrograde, projection-targeted access to mitochondria in distinct neuronal circuits. By combining TOM-Tag with retro-AAV delivery, we enabled molecular and functional characterization of mitochondria based on long-range projection identity rather than marker gene expression alone. This approach expands the versatility of mitochondrial tagging technologies and allows interrogation of mitochondria in subpopulations that are anatomically intermingled but functionally distinct. The validity of this approach was further reinforced by the agreement of translational and proteomic analyses, which uncovered a set of mitochondrial proteins with cell type specificity. Among these, *Cox7a1* stood out as a CSPN-enriched gene at both the transcript and protein levels, uniquely detected in CSPN mitochondrial isolates but absent in CTPNs and bulk cortical mitochondria. RNAscope® confirmed CSPN-restricted expression in situ, highlighting how multi-omic approaches can reveal mitochondrial specializations masked in bulk tissue analysis.

*Cox7a1* encodes a tissue-specific isoform of subunit 7a of cytochrome c oxidase (Complex IV), the terminal component of the electron transport chain (ETC)^46^ . *Cox7a1* is typically characterized as a heart/muscle-specific isoform ^47,48^, where it has been demonstrated to enhance Complex IV activity through its regulation of the assembly and stability of supercomplexes in the ETC^49,50^ . Interestingly, skeletal muscle in ALS is known to exhibit profound oxidative stress, mitochondrial dysfunction, and bioenergetic collapse, suggesting that *Cox7a1*-expressing tissues may share an intrinsic metabolic vulnerability to ALS-related stressors^51^ . Its expression in CSPNs but not in CTPNs or bulk cortical mitochondria suggests that CSPNs may utilize a specialized respirasome to meet their high energetic needs ^50^. Interestingly, recent work by Cheng et al. demonstrated that neuronal deficiency of mitochondrial respiratory Complex IV is sufficient to recapitulate hallmark ALS phenotypes, including motor neuron loss and TDP-43 pathology, underscoring the pathogenic significance of CIV dysfunction in ALS^52^. Additionally, widespread Complex IV deficiency has been reported among patients with sALS^52,53^ . This suggests that a finely tuned Complex IV is critical for motor neuron integrity. The observed COX7A1 specialization may paradoxically render CSPNs more susceptible to mitochondrial dysfunction. Importantly, these findings would not be detectable without a cell-type-resolved, multi-omics approach, highlighting how mitochondrial subunit composition may contribute to neuronal identity and disease risk.

The combined metabolomic and proteomic profiling of CSPN and CTPN mitochondria revealed divergent properties that reflect each cell type’s response to ALS. The selective enrichment of β-oxidation enzymes and cofactors in CTPNs along with increased acyl-carnitines suggest that CTPNs possess a metabolically flexible phenotype capable of leveraging fatty acids to sustain respiration. Additionally, the upregulation of antioxidant enzymes such as SOD1, TXNRD1, and PRDX2 in CTPNs, together with an increased GSH/GSSG ratio, further suggests that their metabolic programming is accompanied by redox buffering, enabling sustained mitochondrial respiration without excessive oxidative damage. This may allow CTPNs to maintain bioenergetic output under stress while limiting oxidative damage, contributing to their relative resilience. In contrast, CSPNs appear metabolically constrained, relying heavily on oxidative phosphorylation and the TCA cycle, with limited capacity for metabolic switching. Under ALS conditions, CSPNs attempt to activate fatty acid metabolic pathways; however, the associated proteomic and metabolomic signatures are less robust, coinciding with evidence of energy depletion and redox imbalance. These observations support a model in which CSPNs operate near their bioenergetic limit in health and fail to adapt under stress. The differential engagement of fatty acid metabolism between these populations adds to a growing body of evidence implicating lipid metabolic dysregulation in ALS pathogenesis^54,55^ . Studies across multiple models and human tissue have shown that altered lipid homeostasis is a common feature of disease, and that enhancing neuronal fatty acid utilization can suppress neurodegeneration and extend survival^56,57^ .

Collectively, our findings provide strong evidence that mitochondrial behavior and plasticity are cell type-specific traits that shape neuronal resilience to ALS-associated stress. The enrichment of respiratory chain components in CSPNs may reflect more exclusive reliance on oxidative phosphorylation to meet their energetic demands making these neurons particularly susceptible to dysfunction under oxidative or metabolic stress. A similar principle has been described in Parkinson’s disease, where dopaminergic neurons in the substantia nigra have been demonstrated to have high oxidative phosphorylation demands and show heightened vulnerability to mitochondrial oxidative and bioenergetic stress^58,59^. Conversely, the metabolic adaptability of CTPNs characterized by broader use of fatty acid oxidation and sustained redox buffering may underlie their resistance. Recent studies have shown that neuronal fatty acid utilization and triglyceride-derived fuel reserves can support synaptic function and memory formation during periods of heightened energetic demand^60–63^ . Our findings extend this framework by suggesting that neurons intrinsically equipped to engage fatty acid pathways may be better positioned to withstand chronic metabolic stress in neurodegenerative disease. These data support a model in which selective neuronal vulnerability in ALS is not solely the result of toxic insult but emerges from intrinsic differences in mitochondrial composition, regulation, and stress response capacity. Beyond ALS, this study offers a generalizable framework for dissecting the subcellular basis of selective vulnerability in the nervous system. Future application of this platform to other degenerative conditions, such as Parkinson’s disease, Alzheimer’s disease, or frontotemporal dementia may uncover valuable insights into mitochondrial programs that shape neuron subtype survival. Finally, our identification of CSPN-specific mitochondrial proteins opens new avenues for developing targeted interventions aimed at bolstering metabolic resilience and preventing cellular distress.

## ACKNOWLEDGEMENTS

This work was supported by NIH/NINDS grants R01NS129902 (E.F.S.) and R21NS133927 (E.F.S.), Simons Foundation SCPAB Fellows to Faculty Fellowship 00003203 and 00017276 (B.A.C.), and Kavli Institute for Neuroscience grant 111630 (B.A.C.). We thank The Rockefeller University Genomics Resource Center, Bioinformatics Resource Center, Electron Microscopy Resource Center, Proteomics Resource Center, Drug Discovery Resource Center and Comparative Bioscience Center. We also thank Dr. Thomas Carroll for additional bioinformatics support, as well as Dr. Tim Kenny for critical discussions, and Jie Xing for technical assistance.

## AUTHOR CONTRIBUTIONS

B.A.C, N.H., and E.F.S. conceived and designed the study. B.A.C., L.Z., B.F., C.P., E.K., H.M., and E.F.S. performed all experiments. B.A.C., E.K., C.P., and E.F.S. analyzed data. B.A.C. wrote the manuscript with comments from N.H. and E.F.S.

## Methods

### Animals

All animal procedures were conducted in accordance with the National Institutes of Health guidelines and approved by the Institutional Animal Care and Use Committee (IACUC) at The Rockefeller University. Tg(Gng7-cre)KH67Gsat mice (MMRRC:031180-UCD; Eric Schmidt), referred to as Gng7-Cre, were used for all experiments and maintained on a C57BL/6J background (Jackson Laboratory, stock #000664, RRID:IMSR_JAX:000664). For ALS model studies, Gng7-Cre mice were crossed with Tg(SOD1^*^G93A)1Gur mice (Jackson Laboratory, stock #004435, RRID:IMSR_JAX:004435). Gng7-Cre::SOD^*^1G93A−/− littermates were used as controls for Gng7-Cre::SOD1^*^G93A+/− experimental animals. SOD1^*^G93A+/− animals exhibiting loss of mobility or inability to feed were euthanized in accordance with humane endpoints defined by IACUC protocol.

Mice were group-housed in a temperature- and humidity-controlled facility under a 12-hour light/dark cycle with ad libitum access to food and water. Both male and female mice were used equally across all experimental conditions.

Seventeen-week-old mice were used for all baseline studies, including vTRAP profiling, TOM-Tag mitochondrial isolation, retrograde viral labeling, histology, and RNAscope®. For SOD1^*^G93A disease-stage analyses, tissue was collected at 17 weeks to represent a symptomatic time point. All spinal cord and intracranial injections were performed between 5 and 7 weeks of age.

### Plasmids and Viral Vectors

For cell type-specific translatome profiling, we used a Cre-dependent vTRAP construct (AAV-FLEX-EGFPL10a), originally developed and validated in our lab^34^. To enable retrograde access to projection neurons, this construct was packaged into the SL1 capsid (AAV2-retro) by the Janelia Viral Tools Facility.

To isolate mitochondria from genetically defined neuronal populations, we generated a Cre-dependent TOM-Tag viral vector. This construct was created by subcloning the mitochondrial outer membrane fusion protein mEmerald-TOMM20 into a DIO pAAV-EF1α backbone previously used for vTRAP constructs. The mEmerald-TOMM20-N-10 plasmid was a gift from Michael Davidson (Addgene plasmid #54282; http://n2t.net/addgene:54282; RRID:Addgene_54282). Homologous recombination was used to insert the TOMM20 fusion coding sequence into the Cre-dependent AAV vector. The final construct (AAV-EF1α-DIO-mEmerald-TOMM20-WPRE-HGHpA) was packaged into the AAV2-retro serotype by the Janelia Viral Tools Facility for retrograde labeling of projection-defined neurons. For in vitro validation of Cre-dependent TOM-Tag expression, we acquired pAAV-Ef1a-mCherry-IRES-Cre as a gift from Karl Deisseroth (Addgene plasmid # 55632 ; http://n2t.net/addgene:55632; RRID:Addgene_55632).

To perform dual retrograde mitochondrial labeling in vivo, we used pAAV-EF1a-mCherry-TOMM20-WPRE-HGHpA, a construct previously generated and described by our lab ^10^ .

### Behavior Testing

#### Rotarod Assessment

Motor coordination and balance were assessed using an accelerating rotarod apparatus. Testing was performed on Gng7-Cre::SOD1^*^G93A+/− mice and their Cre-positive, SOD1G93A−/− littermate controls. Mice were first habituated to the task with a three-day training period beginning at approximately postnatal day 56 (∼P56), three weeks after viral surgery. During training, mice completed two trials per day with a maximum trial length of 180 seconds. The rotarod accelerated linearly from 5 to 18 revolutions per minute (rpm) over 120 seconds, followed by a 60-second hold at 18 rpm.

The day prior to tissue collection, mice underwent a testing session consisting of three trials with the same acceleration protocol. Latency to fall was recorded automatically using an infrared sensor at the base of the apparatus.

### Tissue Collection and Histology

#### Perfusion, fixation, and sectioning

Mice were deeply anesthetized via intraperitoneal injection of ketamine/xylazine (1%/0.1%; 2 mL/kg and 0.2 mL/kg, respectively) and transcardially perfused with ∼40 mL PBS (pH 7.4), followed by ∼30 mL of 4% paraformaldehyde (PFA) in PBS. Brains and spinal cords were post-fixed overnight in 4% PFA at 4 °C with gentle agitation and then cryoprotected in 30% sucrose in PBS.

For immunostaining, tissue was sectioned at 30 μm using a freezing microtome and stored in cryoprotectant solution (25% ethylene glycol, 25% glycerol, 50% PBS) at −20 °C until use. Spinal cords were embedded in Neg-50 medium (Thermo Fisher) on dry ice before sectioning. For RNAscope®, cryosections (14 μm) were cut from OCT-embedded brains using a Leica cryostat and mounted onto Superfrost Plus slides (Thermo Fisher).

#### In situ hybridization (RNAscope®)

RNAscope® fluorescent in situ hybridization was performed using the RNAscope Multiplex Fluorescent Reagent Kit v2 (ACD Bio, #323110) according to the manufacturer’s protocol (#323100-USM). Probes targeting *Vat1l* (#495401), *Serpina3n* (#430191), and *Cox7a1* (#467851) were used to label projection neuron subtypes in the primary motor cortex of wild-type and SOD1G93A brains. Sections were counterstained with DAPI and imaged using a Zeiss LSM 710 confocal microscope under uniform acquisition settings.

#### Immunofluorescence staining

Free-floating 30 μm brain sections were washed in PBS (pH 7.4) to remove cryoprotectant, then blocked in 5% normal donkey serum (Jackson ImmunoResearch) with 0.3% Triton X-100 for 1 h. Primary antibodies were applied overnight at 4 °C in 1% donkey serum and 0.3% Triton X-100. After 3 PBS washes, sections were incubated with species-appropriate fluorescent secondary antibodies for 1 h at room temperature, washed again, and mounted in ProLong Gold Antifade Mountant with DAPI (Thermo Fisher). Imaging was performed using confocal microscopy with identical acquisition settings across samples.

### Stereotaxic and Spinal Cord Injections

Mice were anesthetized with an intraperitoneal injection of ketamine/xylazine (1%/0.1%; 1 mL/kg and 0.1 mL/kg, respectively) in sterile 0.9% saline. For thalamic injections, animals were placed in a stereotaxic frame (Kopf Instruments), and craniotomies were made using a dental drill. Viruses were injected into the ventrolateral thalamus at coordinates: anteroposterior (AP) −1.5 mm, mediolateral (ML) ±1.1 mm, dorsoventral (DV) −3.65 mm from bregma. Incisions were closed using Vetbond (3M). For spinal injections, a midline incision was made at the base of the neck to expose the cervical vertebrae. Back and shoulder muscles were retracted, and the C4 vertebra was partially removed using rongeurs to expose the spinal cord. A small incision was made in the dura, and virus was injected at ML ±0.45 mm, DV −0.95 mm from the spinal surface. In all cases, 0.1–0.15 μL of virus was delivered per site at a rate of 0.1 μL/min using a pulled-glass capillary and Hamilton syringe. After injection, pipettes were left in place for 2 minutes to minimize reflux. Following spinal injections, back and shoulder muscles were sutured, and skin incisions were closed with wound clips. Mice received a single subcutaneous dose of meloxicam sustained release (6 mg/kg) in the interscapular region and were monitored during recovery on a warming pad. Viral expression proceeded for 12 weeks prior to tissue collection.

For dual retrograde mitochondrial labeling, retro-AAV-EF1α-DIO-mEmerald-TOMM20 (TOM-Tag) was injected unilaterally into the VLT and retro-AAV-EF1α-mCherry-TOMM20 into the contralateral side of C4 spinal cord.

For vTRAP translatome profiling, retro-AAV-FLEX-EGFPL10a was injected bilaterally into either the VLT or C4 spinal cord to target corticothalamic or corticospinal projection neurons, respectively.

For mitochondrial tagging, retro-AAV-EF1α-DIO-mEmerald-TOMM20 was injected bilaterally into either the VLT or C4 spinal cord to target corticothalamic or corticospinal projection neurons, respectively.

### Viral Translating Ribosome Affinity Purification (vTRAP)

#### TRAP and RNA Extraction

Affinity purification of EGFP-tagged polysomes was performed 12 weeks post-injection using previously describe method^34,64^ . For each condition, 3–4 biological replicates were collected from dissected M1 cortex. Tissue was homogenized in ice-cold lysis buffer containing: 10 mM HEPES-KOH (pH 7.4), 150 mM KCl, 5 mM MgCl_2_, 0.5 mM DTT, 100 μg/mL cycloheximide, RNasin (Promega), SUPERase-In™ (Life Technologies), and Complete EDTA-free protease inhibitors (Roche). Homogenates were cleared by two-step centrifugation, and EGFP-tagged ribosomes were immunoprecipitated using a 1:1 mixture of anti-GFP monoclonal antibodies (clones 19C8 and 19F7) bound to biotinylated Protein L (Thermo Fisher) on streptavidin-coated magnetic beads overnight (Life Technologies). Bound RNA was purified using the RNeasy Plus Micro Kit (Qiagen).

RNA concentration was measured using a NanoDrop 1000 spectrophotometer, and RNA integrity was assessed with Agilent High Sensitivity RNA ScreenTape (Agilent #5067-5579) on a TapeStation 4200 (Agilent #G2991BA). Only samples with RNA integrity values ≥7.0 were used for downstream sequencing.

#### Library Preparation and Sequencing

RNA-seq libraries were prepared using the Trio™ RNA-Seq Library Preparation Kit (Tecan) according to the manufacturer’s protocol. Libraries were generated from 10 ng of total RNA per sample and analyzed using D1000 ScreenTape on the Agilent TapeStation. Sequencing was performed at The Rockefeller University Genomics Resource Center using an Illumina NextSeq 500 system (paired-end, 75 bp, high-output mode).

### Sequencing analyses

Sequence and transcript coordinates for mouse mm10 UCSC genome and gene models were retrieved from the Bioconductor Bsgenome.Mmusculus.UCSC.mm10 (version 1.4.0) and TxDb.Mmusculus.UCSC.mm10.knownGene (version 3.4.0) Bioconductor libraries respectively. RNA-seq reads are aligned to the genome using the subjunc method (version 1.30.6) in Rsubread (Liao et al., 2013) and exported as bigWigs normalized to reads per million using the rtracklayer package (version 1.40.6). Counts in genes were generated using featureCounts within the Rsubread package (version 1.30.6) with default settings and the TxDb.Mmusculus.UCSC.mm10.knownGene gene models. Normalization and rlog transformation of raw read counts in genes were performed using DESeq2 (version 1.44.0) and between sample variability was assessed with hierarchical clustering and heat maps of between sample distances implemented in the pheatmap R package (1.0.13). Differential expression analysis on gene counts was performed using all default parameters in DESeq2; in RStudio (version 2024.4.2.764; https://www.rstudio.com/), with R (version 4.4.1; https://www.r-project.org). Gene ontology enrichment analyses were performed using clusterProfiler package (version 4.16.6) with default parameters, with genes with mean counts per million >100 being used for these analyses.

#### In Vitro Transfection and Mitochondrial Labeling

HEK293T cells were cultured in DMEM (Gibco) supplemented with 10% fetal bovine serum (FBS; Gibco) and 1% penicillin-streptomycin (Gibco) and maintained at 37 °C in a humidified incubator with 5% CO_2_. Cells were seeded onto glass-bottom 96-well dishes at ∼60% confluency and transfected the next day using Lipofectamine 3000 (Thermo Fisher) according to the manufacturer’s protocol.

For TOM-Tag expression, cells were co-transfected with pAAV-EF1α-DIO-mEmerald-TOMM20 and pAAV-EF1α-mCherry-IRES-Cre (Addgene plasmid #55632; RRID:Addgene_55632) at a 1:1 ratio. Twenty-four hours post-transfection, cells were incubated with 100 nM MitoTracker™ Deep Red FM (Thermo Fisher, #M22426) diluted in prewarmed culture medium for 15 min at 37 °C. Cells were then washed 3× with PBS and fixed in 4% paraformaldehyde for 15 min at room temperature. Cells were then washed 3× with PBS containing DAPI and imaged by confocal microscopy.

### Mitochondrial Isolation

#### Antibody–Bead Conjugation

All steps were performed on ice or at 4 °C unless otherwise indicated. Streptavidin MyOne T1 Dynabeads (Thermo Fisher) were used to capture anti-EGFP antibodies via biotinylated Protein L. For each sample, 150 μL of bead solution was transferred to a 1.5-mL microcentrifuge tube and placed on a magnetic stand at room temperature for 2 min to collect beads. After removing the supernatant, beads were washed once with 1 mL PBS by gentle inversion and returned to the magnet. Beads were resuspended in 880 μL PBS, and 120 μL of freshly reconstituted biotinylated Protein L (1 mg/mL; Thermo Fisher) was added. The mixture was incubated for 35 min at room temperature on a tube rotator. Beads were then washed 5× with PBS containing 3% nuclease-free BSA (Ambion), followed by a final resuspension in 500 μL 0.15 M KCl wash buffer (150 mM KCl, 10 mM HEPES (pH 7.4), 5 mM MgCl_2_, 1% NP-40). Mouse monoclonal anti-EGFP antibodies (clones 19C8 and 19F7) were pre-spun at 4 °C and added at 50 μg each (total 100 μg) per sample. Beads were incubated with antibody for 1 h at room temperature on a slow tube rotator, washed 3× with ice-cold mitochondrial isolation buffer (MIB, 37 mM KCl, 3 mM KH_2_PO_4_, 2.5 mM MgCl_2_, 10 mM HEPES, 1 mM EDTA; pH adjusted to 7.4 with KOH.), and resuspended in 200 μL MIB containing 1% fatty acid–free BSA and protease inhibitors.

#### Mitochondrial Immunopurification

For each sample, ∼50 mg of dissected M1 cortex (25 mg per hemisphere) was homogenized in 2 mL MIB with BSA using a Dounce homogenizer (6 strokes with loose pestle A, then 16 strokes with tight pestle B). Homogenate was centrifuged at 1,300 × g for 3 min at 4 °C. The supernatant was transferred to a pre-chilled 15-mL tube. The pellet was resuspended in 2 mL MIB with BSA, homogenized again with pestle B (6 strokes), and centrifuged again at 1,300 × g for 3 min. Both supernatants were combined and incubated with prewashed anti-GFP magnetic beads for 1 h at 4 °C on a tube rotator. Whole tissue lysate and post-spin input fraction was saved. Beads were collected with a magnet, and the unbound fraction was saved. Bound mitochondria were gently resuspended in 1 mL MIB with BSA and transferred to a clean 1.5-mL tube to minimize background. Beads were washed 3× with 1 mL ice-cold MIB without BSA by gentle inversion. The final immunopurified (IP) mitochondrial fraction was processed based on downstream application.

#### Transmission Electron Microscopy

Immunopurified mitochondria along with an aliquot of the input fraction were embedded in 2% low-melting-point agarose prepared in 0.1 M sodium cacodylate buffer, pH 7.2 and fixed in 2% glutaraldehyde, 4% PFA, 2 mM CaCl2 in 0.1 M sodium cacodylate buffer, pH 7.2, for 2h at room temperature with gentle agitation, post-fixed in 1% osmium tetroxide, en bloc stained with uranyl acetate, dehydrated through a graded acetone series and processed for Eponate 12 epoxy resin (Ted Pella Inc., USA) embedding.

Ultrathin sections (∼65 nm) were cut and contrasted with 1% aqueous uranyl acetate followed by Reynolds’ lead citrate. Contrasted sections were imaged in a Tecnai 12 transmission electron microscope (FEI, Hillsboro, OR) operated at an accelerating voltage of 120 kV. Digital images were acquired using an AMT BioSprint29 CCD camera (AMT, Woburn, MA).

#### Western Blotting

Immunopurified mitochondrial fractions were lysed directly on magnetic beads in 30 μL RIPA buffer. Samples were incubated on ice for 30 min, then placed on magnet to remove beads. The remaining solution was centrifuged at 21,000 × g for 10 min at 4 °C. BCA was performed to quantify total protein. Protein lysates were mixed with 4× LDS sample buffer (Invitrogen) and 10× reducing agent and heated at 95 °C for 5 min. Samples were run on a 4–12% Bis-Tris precast polyacrylamide gels (Invitrogen) at 200 V for 50 min in 1× MOPS SDS running buffer. Proteins were transferred to PVDF membranes using iBlot 2 dry transfer system using P0 setting (Thermo). Membranes were blocked in LI-COR blocking buffer for 1 h at room temperature, then incubated with primary antibodies overnight at 4 °C in antibody solution (Licor) (typical dilutions 1:500–1:1000, per antibody datasheet). After washing 3× for 5 min in TBST, membranes were incubated with fluorescent secondary antibodies (LI-COR, 1:10,000) for 1 h at room temperature, followed by another 3× TBST wash. Blots were imaged using a LI-COR Odyssey imaging system. Membranes were stripped using Newblot PVDF stripping buffer following manufactured protocol (Licor).

#### Differential Centrifugation for Crude Cortical Mitochondria Isolation

All procedures were conducted on ice or at 4 °C unless otherwise noted. Approximately 50 mg of M1 cortical tissue (25 mg per hemisphere) was dissected, rinsed three times in ice-cold PBS, and homogenized using the MIB buffer and Dounce homogenization procedure described above for mitochondrial immunoprecipitation. The homogenate was centrifuged at 700 × g for 10 min at 4 °C to pellet nuclei and debris. The supernatant was transferred to new pre-chilled tubes, and the low-speed centrifugation step was repeated once more to ensure complete removal of cellular debris. The resulting supernatant was then centrifuged at 10,000 × g for 15 min at 4 °C to pellet the crude mitochondrial fraction. The pellet was resuspended in buffer containing 0.02% digitonin to improve outer membrane purity and centrifuged again at 10,000 × g for 15 min at 4 °C. The final mitochondrial pellet was resuspended in appropriate assay buffer depending on downstream applications.

#### Mitochondrial Respiration Assay (Seahorse XF96 Mito Stress Test)

Mitochondrial oxygen consumption rate (OCR) was measured using the Seahorse XF96 Extracellular Flux Analyzer (Agilent) and the XF Cell Mito Stress Test adapted for isolated mitochondria^65^ . Assays were performed in 96-well Seahorse plates using mitochondria immunopurified via TOM-Tag. The day prior to the assay, sensor cartridges were hydrated by adding 200 μL sterile water and incubated overnight at 37 °C in a non-CO_2_ incubator. XF Calibrant (Agilent) was also prewarmed overnight at 37 °C.

On the day of the assay, isolated mitochondria were diluted in mitochondrial assay solution (MAS; 220 mM mannitol, 70 mM sucrose, 10 mM KH_2_PO_4_, 5 mM MgCl_2_, 2 mM HEPES, 1 mM EGTA, 0.2% fatty acid–free BSA, pH 7.2 with KOH) supplemented with 10 mM succinate + 2 μM rotenone (Complex II). Protein concentration was determined by BCA assay (Thermo Fisher), and mitochondria were diluted to 1 μg/μL.

Ten microliters of mitochondrial suspension (10 μg/well) was pipetted into each well of a Seahorse XF96 microplate, along with 10 μL of MAS in blank wells. Plates were centrifuged at 2,000 × g for 20 min at 4 °C in a swinging-bucket rotor. Wells were visually inspected to confirm adherence of mitochondrial pellets. Subsequently, 170 μL of prewarmed MAS containing substrates and 4 mM ADP was added to each well for a final volume of 180 μL. Plates were incubated at 37 °C in a non-CO_2_ incubator for 10 min prior to starting the assay. Mitochondrial respiration was measured in response to sequential injection of the following inhibitors and uncouplers (final concentrations): 5 μM oligomycin (ATP synthase inhibitor), 4 μM FCCP (uncoupler), and 10 μM antimycin A + 2 μM rotenone (Complex III/I inhibitors). All reagents were prepared in MAS without BSA and loaded into injection ports of the hydrated sensor cartridge as 10× stocks. The XF96 run was initiated following sensor calibration using Wave software. Data were analyzed using Wave (Agilent) and Prism (GraphPad) software.

#### Metabolomics

Polar metabolites were extracted from immunopurified mitochondrial fractions by adding 50 μL of ice-cold methanol:water (80:20, v/v) containing 1 μM 13C, 15N-labeled amino acids (MSK-A2-1.2, Cambridge Isotope Laboratories, Inc.). After rotating samples for 10 min at 4 °C, magnetic beads were separated, and the supernatant was transferred to a new tube, centrifuged at 20,000 × g for 10 min at 4 °C. A pooled quality control (QC) sample, composed of aliquots from each sample, was run throughout the LC–MS sequence to monitor instrument variability such as mass error, chromatographic shift, and internal standard signal. All samples were analyzed using an IQ-X Orbitrap mass spectrometer coupled to a Vanquish UHPLC system (Thermo Fisher Scientific). For chromatographic separation, samples were loaded on a SeQuant ZIC-pHILIC column (150 × 2.1 mm, 5-μm polymeric) and a SeQuant ZIC-pHILIC Guard Kit (20 × 2.1 mm) at 40 °C and eluted with a solvent system composed of mobile phase A (20 mM ammonium carbonate and 0.1% ammonium hydroxide in water at pH 9.3) and mobile phase B (100% acetonitrile). The injection volume was set to 5 μL, and samples were maintained at 4 °C. The gradient (v/v) used was as follows: 0–22 min, linear gradient from 90% to 40% B; 22–24 min, held at 40% B; 24–24.1 min, returned to 90% B; 24.1–30 min, equilibrated at 90% B at a flow rate of 150 μL/min. Mass spectrometry was performed in polarity-switching mode with spray voltages of 3.0 kV (positive) and 2.5 kV (negative), sheath gas at 40 arbitrary units (AU), auxiliary gas at 15 AU, and both ion transfer tube and vaporizer temperatures set to 300 °C. Data were acquired at a resolution of 120,000 with an automatic gain control (AGC) target of 1.2 × 10^5^, maximum injection time of 246 ms, and two mass ranges: 55–275 and 265– 1000 m/z. External mass calibration was performed weekly using Pierce FlexMix Calibration Solution (Thermo Fisher Scientific), and internal calibration was enabled during acquisition using the EASY-IC system. Relative quantitation of targeted metabolites was performed using Skyline Daily (version 24.1.1.284).

#### Data analysis

Data analysis was conducted in R (v4.4.1). Features were filtered to retain those with a biological-to-blank fold change ≥5 and present in ≥80% of QC samples and ≥70% of biological samples. Internal standard (IS) normalization was performed using a composite IS calculated as the median of all IS with QC coefficient of variation (CV) ≤15% and robust CV ≤30%; blank samples were excluded from normalization. Run-order drift correction was applied using QC-based locally estimated scatterplot smoothing (LOESS) regression (qcrlscR package version 0.1.3, divide method, polynomial degree 2, with optimized span). Features with QC CV ≤30% after drift correction were retained for further analysis. Residual batch effects were removed using the EigenMS algorithm (malbacR package version 0.1.0) applied to log_2_-transformed data from biological samples only, while preserving the biological class factor.

Z-scores were calculated and TCA Cycle pathway diagram generated using MetaboMAPS (https://metabomaps.brenda-enzymes.org/)^42^ . Metabolite ratios were computed on linear-scale values. Group differences were assessed using one-way ANOVA across the four experimental conditions (CSPN_WT, CSPN_SOD, CTPN_WT, CTPN_SOD) in MetaboAnalyst (www.metaboanalyst.ca). Normalized, log2-transformed intensities were uploaded without additional transformation or scaling. P values were adjusted for multiple testing using the Benjamini–Hochberg false discovery rate (FDR) procedure, and metabolites with FDR < 0.05 were considered statistically significant.

Metabolic pathway enrichment analysis was performed using MetaboAnalyst 6.0 with over-representation analysis (ORA) against Small Molecule Pathway Database (SMPDB) pathway sets. All significant metabolites identified by ANOVA FDR were used as input for ORA. Enrichment ratios were calculated as the number of observed hits divided by the number of expected hits, with statistical significance determined using Benjamini–Hochberg correction. Data visualizations included per-sample boxplots of log_10_-transformed intensities, principal component analysis (PCA) of mean-centered and unit-variance scaled log-transformed data, and heatmaps of per-metabolite z-scores clustered by experimental condition.

#### Proteomics and Mass Spectrometry

Mitochondria immunoprecipitated via TOM-Tag were lysed directly on magnetic beads using 25 μL of EasyPep lysis buffer (Thermo Fisher Scientific) and incubated on ice for 30 min. Following incubation, samples were magnetized to pellet the beads, and the cleared lysate was transferred to fresh tubes. Lysates were further clarified by centrifugation at 21,000 × g for 10 minutes at 4 °C. A total of 10 μg of protein from each sample was processed for a quantitative proteomics profiling. In short: Samples were reduced (10mM DTT) and alkylated (30mM iodoacetamide) and ice-cold acetone precipitated overnight. Pellets were collected by centrifugation, air-dried, and resuspended in digestion buffer. Proteins were digested with trypsin (Promega) overnight at 37 °C, and the reaction was halted by addition of neat trifluoroacetic acid (Fisher Scientific). A portion of each digested crude sample was pooled and subjected to high pH reversed-phase fractionation (Thermo Fisher Scientific; P/N 84868), generating 8 fractions. Peptides were cleaned by solid phase extraction (SPE) to remove salts and other impurities prior to LC–MS/MS analysis.

Peptides were analyzed by data-independent acquisition (DIA) using a Thermo Scientific Q Exactive HF Orbitrap mass spectrometer coupled to a Dionex 3000 with a trap column setup. Individual samples and pooled fractions were separated on a 100μm/12-cm pulled emitter C18 reversed-phase column (Nikkyo Technos; P/ N NTCC-360/100-3-125) using a 60-minute analytical gradient (BufferA:Buffer B composition being 1% acetonitrile,0.1% formic acid: 80% acetonitrile, 0.1% formic acid, with a %B gradient of 1 – 38% at 900 nl/min).The fractionated pooled sample was analyzed using data-dependent acquisition (DDA). The instrument was operated in high-resolution/high-mass-accuracy DIA or DDA mode with optimized acquisition parameters.

#### Data Analysis

DDA data files were queried against a Uniprot Mus Musculus database using Proteome Discover v.3.0/SequestHT (Thermo Fisher Scientific). Matched peptides were used to generate a peptide spectral library. DIA data were analyzed by Spectronaut (v.19; Biognosys) using library-assisted Direct-DIA. All DIA data were searched against a Uniprot Mus Musculus database and the DDA generated peptide spectral library. Cross-run normalization was enabled. Protein quantification and differential expression analyses were performed using the MS-DAP (Mass Spectrometry Downstream Analysis Pipeline) package^66^ . This included normalization, statistical modeling, and visualization of protein abundance across conditions. Mitochondrial proteins were annotated using MitoCarta 3.0, and Gene Ontology enrichment analyses were performed on significant differentially expressed proteins using statistical thresholds (q < 0.05, |log_2_FC| ≥ 1 unless otherwise stated).

#### Confocal Imaging

Confocal fluorescence imaging was performed using a Zeiss LSM 710 laser scanning confocal microscope equipped with standard excitation lasers (405, 488, 561, and 633 nm) and a 10x, 20x, or 63x oil-immersion objective. Images of fixed tissue sections and cell culture samples were acquired using Zen software (Zeiss), with the same acquisition parameters applied across experimental groups within each experiment to allow for comparative analysis.

## References

1. Rowland, L. P. & Shneider, N. A. Amyotrophic Lateral Sclerosis. N. Engl. J. Med. 344, 1688– 1700 (2001).

2. Feldman, E. L. et al. Amyotrophic lateral sclerosis. Lancet 400, 1363–1380 (2022).

3. Bos, M. A. J. van den, Geevasinga, N., Higashihara, M., Menon, P. & Vucic, S. Pathophysiology and Diagnosis of ALS: Insights from Advances in Neurophysiological Techniques. Int. J. Mol. Sci. 20, 2818 (2019).

4. Marques, C., Burg, T., Scekic-Zahirovic, J., Fischer, M. & Rouaux, C. Upper and Lower Motor Neuron Degenerations Are Somatotopically Related and Temporally Ordered in the Sod1 Mouse Model of Amyotrophic Lateral Sclerosis. Brain Sci. 11, 369 (2021).

5. Zang, D. W. & Cheema, S. S. Degeneration of corticospinal and bulbospinal systems in the superoxide dismutase 1G93A G1H transgenic mouse model of familial amyotrophic lateral sclerosis. Neurosci. Lett. 332, 99–102 (2002).

6. Liu, Y. et al. C9orf72 BAC Mouse Model with Motor Deficits and Neurodegenerative Features of ALS/FTD. Neuron 90, 521–534 (2016).

7. Özdinler, P. H. et al. Corticospinal Motor Neurons and Related Subcerebral Projection Neurons Undergo Early and Specific Neurodegeneration in hSOD1G93A Transgenic ALS Mice. J. Neurosci. 31, 4166–4177 (2011).

8. Fil, D. et al. Mutant Profilin1 transgenic mice recapitulate cardinal features of motor neuron disease. Hum. Mol. Genet. 26, 686–701 (2017).

9. Yasvoina, M. V. et al. eGFP Expression under UCHL1 Promoter Genetically Labels Corticospinal Motor Neurons and a Subpopulation of Degeneration-Resistant Spinal Motor Neurons in an ALS Mouse Model. J. Neurosci. 33, 7890–7904 (2013).

10. Moya, M. V. et al. Unique molecular features and cellular responses differentiate two populations of motor cortical layer 5b neurons in a preclinical model of ALS. Cell Rep. 38, 110556 (2022).

11. Rossner, M. J. et al. Global Transcriptome Analysis of Genetically Identified Neurons in the Adult Cortex. J. Neurosci. 26, 9956–9966 (2006).

12. Nolan, M., Scott, C., Hof, Patrick. R. & Ansorge, O. Betz cells of the primary motor cortex. J. Comp. Neurol. 532, e25567 (2024).

13. Le Masson, G., Przedborski, S. & Abbott, L. F. A Computational Model of Motor Neuron Degeneration. Neuron 83, 975–988 (2014).

14. Monzel, A. S., Enríquez, J. A. & Picard, M. Multifaceted mitochondria: moving mitochondrial science beyond function and dysfunction. Nat. Metab. 5, 546–562 (2023).

15. Pekkurnaz, G. & Wang, X. Mitochondrial heterogeneity and homeostasis through the lens of a neuron. Nature Metabolism 4, 802–812 (2022).

16. Johnson, D. T., Harris, R. A., Blair, P. V. & Balaban, R. S. Functional consequences of mitochondrial proteome heterogeneity. Am. J. Physiol.-Cell Physiol. 292, C698–C707 (2007).

17. Johnson, D. T. et al. Tissue heterogeneity of the mammalian mitochondrial proteome. American Journal of Physiology-Cell Physiology 292, C689–C697 (2007).

18. Pagliarini, D. J. et al. A Mitochondrial Protein Compendium Elucidates Complex I Disease Biology. Cell 134, 112–123 (2008).

19. McLaughlin, K. L. et al. Novel approach to quantify mitochondrial content and intrinsic bioenergetic efficiency across organs. Sci. Rep. 10, 17599 (2020).

20. Benard, G. et al. Physiological diversity of mitochondrial oxidative phosphorylation. Am. J. Physiol.-Cell Physiol. 291, C1172–C1182 (2006).

21. Petersen, M. H. et al. Functional Differences between Synaptic Mitochondria from the Striatum and the Cerebral Cortex. Neuroscience 406, 432–443 (2019).

22. Faitg, J. et al. 3D neuronal mitochondrial morphology in axons, dendrites, and somata of the aging mouse hippocampus. Cell Rep. 36, 109509 (2021).

23. Graham, L. C. et al. Proteomic profiling of neuronal mitochondria reveals modulators of synaptic architecture. Mol. Neurodegener. 12, 77 (2017).

24. Völgyi, K. et al. Synaptic mitochondria: A brain mitochondria cluster with a specific proteome. J. Proteom. 120, 142–157 (2015).

25. Mishra, J. et al. Differential Ca2+ handling by isolated synaptic and non-synaptic mitochondria: roles of Ca2+ buffering and efflux. Front. Synaptic Neurosci. 17, 1562065 (2025).

26. Fecher, C. et al. Cell-type-specific profiling of brain mitochondria reveals functional and molecular diversity. Nat. Neurosci. 22, 1731–1742 (2019).

27. Lee, J., Pye, N., Ellis, L., Vos, K. D. & Mortiboys, H. Evidence of mitochondrial dysfunction in ALS and methods for measuring in model systems. Int. Rev. Neurobiol. 176, 269–325 (2024).

28. Vandoorne, T., Bock, K. D. & Bosch, L. V. D. Energy metabolism in ALS: an underappreciated opportunity? Acta Neuropathol. 135, 489–509 (2018).

29. Gautam, M., Gunay, A., Chandel, N. S. & Ozdinler, P. H. Mitochondrial dysregulation occurs early in ALS motor cortex with TDP-43 pathology and suggests maintaining NAD+ balance as a therapeutic strategy. Sci. Rep. 12, 4287 (2022).

30. Schweingruber, C. et al. Single-cell RNA-sequencing reveals early mitochondrial dysfunction unique to motor neurons shared across FUS- and TARDBP-ALS. Nat. Commun. 16, 4633 (2025).

31. Gautam, M., Xie, E. F., Kocak, N. & Ozdinler, P. H. Mitoautophagy: A Unique Self-Destructive Path Mitochondria of Upper Motor Neurons With TDP-43 Pathology Take, Very Early in ALS. Front. Cell. Neurosci. 13, 489 (2019).

32. Gurney, M. E. et al. Motor Neuron Degeneration in Mice that Express a Human Cu,Zn Superoxide Dismutase Mutation. Science 264, 1772–1775 (1994).

33. Economo, M. N. et al. Distinct descending motor cortex pathways and their roles in movement. Nature 563, 79–84 (2018).

34. Nectow, A. R. et al. Rapid Molecular Profiling of Defined Cell Types Using Viral TRAP. Cell Rep. 19, 655–667 (2017).

35. Tervo, D. G. R. et al. A Designer AAV Variant Permits Efficient Retrograde Access to Projection Neurons. Neuron 92, 372–382 (2016).

36. Bannwarth, S. et al. A mitochondrial origin for frontotemporal dementia and amyotrophic lateral sclerosis through CHCHD10 involvement. Brain 137, 2329–2345 (2014).

37. Johnson, J. O. et al. Mutations in the CHCHD10 gene are a common cause of familial amyotrophic lateral sclerosis. Brain 137, e311–e311 (2014).

38. Alves, C. J. et al. Early motor and electrophysiological changes in transgenic mouse model of amyotrophic lateral sclerosis and gender differences on clinical outcome. Brain Res. 1394, 90–104 (2011).

39. Mancuso, R., Oliván, S., Osta, R. & Navarro, X. Evolution of gait abnormalities in SOD1G93A transgenic mice. Brain Res. 1406, 65–73 (2011).

40. Bayraktar, E. C. et al. MITO-Tag Mice enable rapid isolation and multimodal profiling of mitochondria from specific cell types in vivo. Proc. Natl. Acad. Sci. 116, 303–312 (2019).

41. Lewis, S. C., Uchiyama, L. F. & Nunnari, J. ER-mitochondria contacts couple mtDNA synthesis with mitochondrial division in human cells. Science 353, aaf5549 (2016).

42. Koblitz, J., Schomburg, D. & Neumann-Schaal, M. MetaboMAPS: Pathway sharing and multi-omics data visualization in metabolic context. F1000Research 9, 288 (2020).

43. Mello, N. P. de, Fecher, C., Pastor, A. M., Perocchi, F. & Misgeld, T. Ex vivo immunocapture and functional characterization of cell-type-specific mitochondria using MitoTag mice. Nat. Protoc. 18, 2181–2220 (2023).

44. Pastor, A. M. et al. Neuronal compartmentalization results in impoverished axonal mitochondria. (2025) doi:10.1101/2025.10.27.684882.

45. Gella, A. et al. Mitochondrial Proteome of Affected Glutamatergic Neurons in a Mouse Model of Leigh Syndrome. Front. Cell Dev. Biol. 8, 660 (2020).

46. Pham, L. et al. Regulation of mitochondrial oxidative phosphorylation through tight control of cytochrome c oxidase in health and disease – Implications for ischemia/reperfusion injury, inflammatory diseases, diabetes, and cancer. Redox Biol. 78, 103426 (2024).

47. Vercellino, I. & Sazanov, L. A. Structure and assembly of the mammalian mitochondrial supercomplex CIII2CIV. Nature 598, 364–367 (2021).

48. Jaradat, S. A., Ko, M. S. H. & Grossman, L. I. Tissue-Specific Expression and Mapping of theCox7ahGene in Mouse. Genomics 49, 363–370 (1998).

49. García-Poyatos, C. et al. Cox7a1 controls skeletal muscle physiology and heart regeneration through complex IV dimerization. Dev. Cell 59, 1824-1841.e10 (2024).

50. Fernández-Vizarra, E. et al. Two independent respiratory chains adapt OXPHOS performance to glycolytic switch. Cell Metab. 34, 1792-1808.e6 (2022).

51. Loeffler, J., Picchiarelli, G., Dupuis, L. & Aguilar, J. G. D. The Role of Skeletal Muscle in Amyotrophic Lateral Sclerosis. Brain Pathol. 26, 227–236 (2016).

52. Cheng, M. et al. Mitochondrial respiratory complex IV deficiency recapitulates amyotrophic lateral sclerosis. Nat. Neurosci. 28, 748–756 (2025).

53. Crugnola, V. et al. Mitochondrial Respiratory Chain Dysfunction in Muscle From Patients With Amyotrophic Lateral Sclerosis. Arch. Neurol. 67, 849–854 (2010).

54. Lee, H. et al. Multi-omic analysis of selectively vulnerable motor neuron subtypes implicates altered lipid metabolism in ALS. Nat. Neurosci. 24, 1673–1685 (2021).

55. Hanrieder, J. & Ewing, A. G. Spatial Elucidation of Spinal Cord Lipid- and Metabolite-Regulations in Amyotrophic Lateral Sclerosis. Sci. Rep. 4, 5266 (2014).

56. Guo, K. et al. Longitudinal Metabolomics in Amyotrophic Lateral Sclerosis Implicates Impaired Lipid Metabolism. Ann. Neurol. 98, 19–34 (2025).

57. Giblin, A. et al. Neuronal polyunsaturated fatty acids are protective in ALS/FTD. Nat. Neurosci. 28, 737–747 (2025).

58. Pacelli, C. et al. Elevated Mitochondrial Bioenergetics and Axonal Arborization Size Are Key Contributors to the Vulnerability of Dopamine Neurons. Curr. Biol. 25, 2349–2360 (2015).

59. Zampese, E. et al. Ca2+ channels couple spiking to mitochondrial metabolism in substantia nigra dopaminergic neurons. Sci. Adv. 8, eabp8701 (2022).

60. Pavlowsky, A. et al. Neuronal fatty acid oxidation fuels memory after intensive learning in Drosophila. Nat. Metab. 7, 2438–2450 (2025).

61. Kumar, M. et al. Triglycerides are an important fuel reserve for synapse function in the brain. Nat. Metab. 7, 1392–1403 (2025).

62. Greda, A. K. et al. Interaction of sortilin with apolipoprotein E3 enables neurons to use long-chain fatty acids as alternative metabolic fuel. Nat. Metab. 7, 2346–2365 (2025).

63. Saber, S. H. et al. DDHD2 provides a flux of saturated fatty acids for neuronal energy and function. Nat. Metab. 7, 2117–2141 (2025).

64. Heiman, M., Kulicke, R., Fenster, R. J., Greengard, P. & Heintz, N. Cell type–specific mRNA purification by translating ribosome affinity purification (TRAP). Nat. Protoc. 9, 1282–1291 (2014).

65. Rogers, G. W. et al. High Throughput Microplate Respiratory Measurements Using Minimal Quantities Of Isolated Mitochondria. PLoS ONE 6, e21746 (2011).

66. Koopmans, F., Li, K. W., Klaassen, R. V. & Smit, A. B. MS-DAP Platform for Downstream Data Analysis of Label-Free Proteomics Uncovers Optimal Workflows in Benchmark Data Sets and Increased Sensitivity in Analysis of Alzheimer’s Biomarker Data. J. Proteome Res. 22, 374– 386 (2023).

