## Supplemental Figures for "Divergent mitochondrial and metabolic adaptations shape selective vulnerability in ALS"

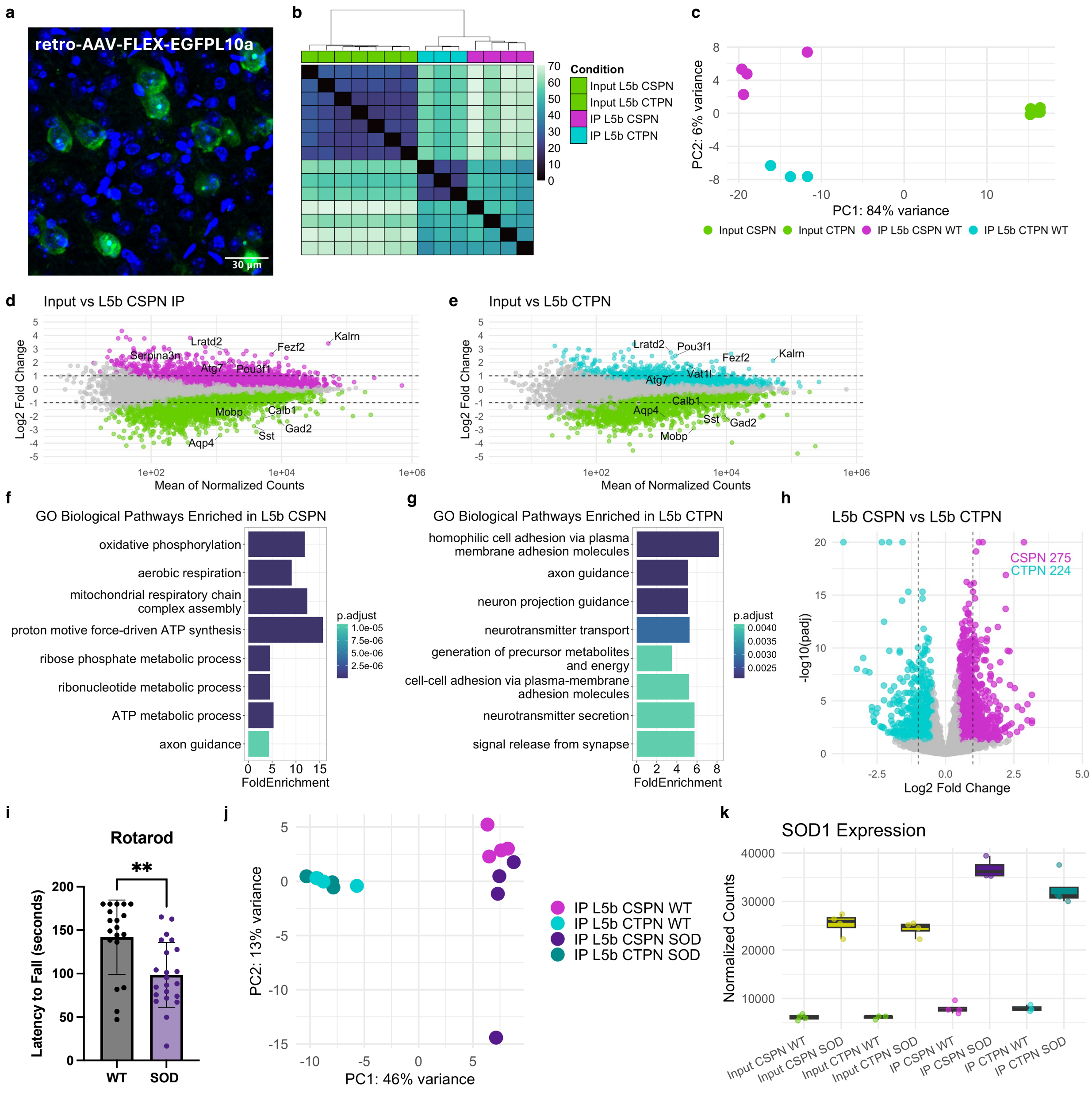


**Supplementary Figure 1. Validation of selective vulnerability, vTRAP targeting, and translatome profiling of CSPNs and CTPNs. a,** High magnification image of anti-GFP immunofluorescence following retro-AAV-FLEX-EGFPL10a injection into C4–C5 (CSPNs) in Gng7-Cre mice shows cell body–restricted cytoplasmic labeling of the transgene. **b,** Hierarchical clustering of all RNA-seq samples based on normalized expression of all genes separates input, CSPN, and CTPN vTRAP samples into distinct clusters. **c)** PCA plot of the top 500 most variable genes across samples further separates CSPN IPs (magenta), CTPN IPs (cyan), and inputs from both cell types (green) into distinct groups. **d–e**, MA plots showing differential expression between CSPN IP (e) and CTPN IP (f) TRAP samples relative to whole tissue input. DEGs (padj < 0.05, baseMean > 100) enriched in CSPNs (magenta, e), CTPNs (cyan, f) are indicated. Dotted horizontal lines indicate $\log_2$ fold change of ±1. Green points represent genes enriched in input and thus depleted from the IPs. **f–g,** Top GO terms for genes enriched in CSPN IPs (g) and CTPN IPs (h) over input. **h,** Volcano plot identifying DEGs between CSPN and CTPN TRAP IPs. Genes significantly enriched (padj <0.05) in CSPNs (magenta) or CTPNs (cyan) are indicated Dotted vertical lines denote $\log_2$ fold change of ±1. Number of DEGs for each cell type is stated. **i)** Bar graph showing rotarod performance (Mean $\pm$ SEM) of Gng7-Cre::SOD1G93A mice and WT littermates at postnatal day 120 (P120), measured as latency to fall (seconds). Each point represents an individual animal. **p < 0.001, unpaired two-tailed Student's t-test. **j)** PCA plot of vTRAP samples from Gng7-Cre WT (light colors) and SOD1*G93A (dark) mice shows separation between CSPNs (magenta) and CTPNs (cyan), with additional clustering by mutant SOD1 expression in CSPNs only. **k)** Box plot showing normalized expression of the human SOD1 transgene in all v TRAP samples from Gng7-Cre::SOD1G93A mice.

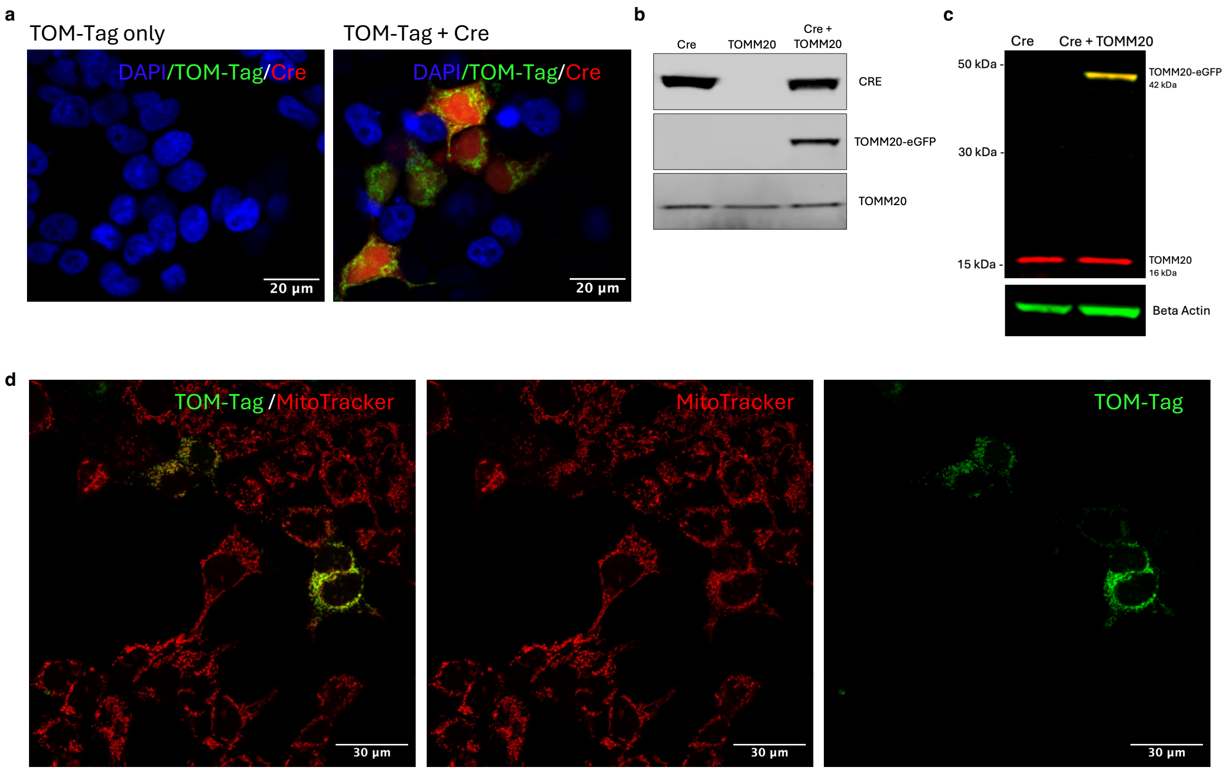


**Supplementary Figure 2. In vitro validation of TOM-Tag construct in HEK293T cells. a**, Confocal images of HEK293T cells transfected with TOM-Tag (mEmerald-TOMM20) alone or co-transfected with an mCherry and Cre co-expressing plasmid. **b**, Western blot of HEK293T lysates transfected with Cre only, TOM-Tag only, or TOM-Tag plus Cre. Blots were probed for Cre and mEmerald-TOMM20. **c**, Western blot comparing TOM-Tag expression (mEmerald-TOMM20) in cells transfected with Cre alone or TOM-Tag plus Cre. β-Actin is shown as a loading control. **d**, High magnification confocal images of HEK293T cells co-transfected with TOM-Tag and Cre, stained with MitoTrackerRed.

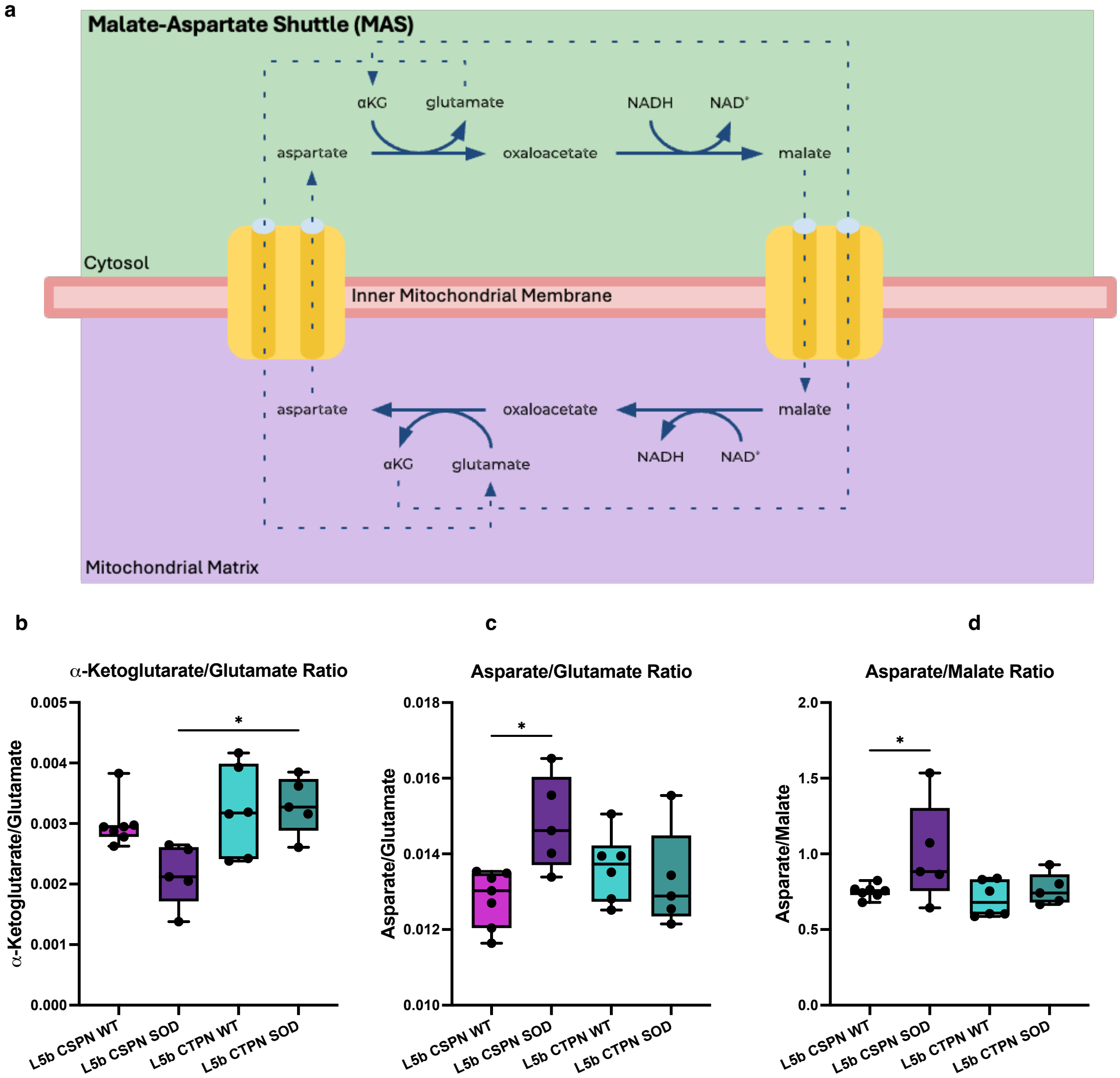


**Supplementary Figure 3. TCA intermediate ratios and redox status in CSPN and CTPN mitochondria. a**, Schematic of the malate–aspartate shuttle illustrating the mitochondrial exchange of key metabolites, including malate, aspartate, α-ketoglutarate, and glutamate, across the inner mitochondrial membrane. Metabolites quantified in this study are highlighted. **b**, Box plot showing the ratio of α-ketoglutarate to glutamate in mitochondrial fractions isolated from CSPNs and CTPNs under WT and SOD1*G93A conditions. **c**, Box plot of aspartate to glutamate ratio across the same conditions as b. **d**, Box plot of aspartate to malate ratio in each condition. Each point represents an individual biological replicate. Ratios were calculated from normalized LC–MS metabolite abundances in TOM-Tag–immunoprecipitated mitochondria. Statistical significance was assessed using one-way ANOVA with multiple comparisons. *p = 0.01.

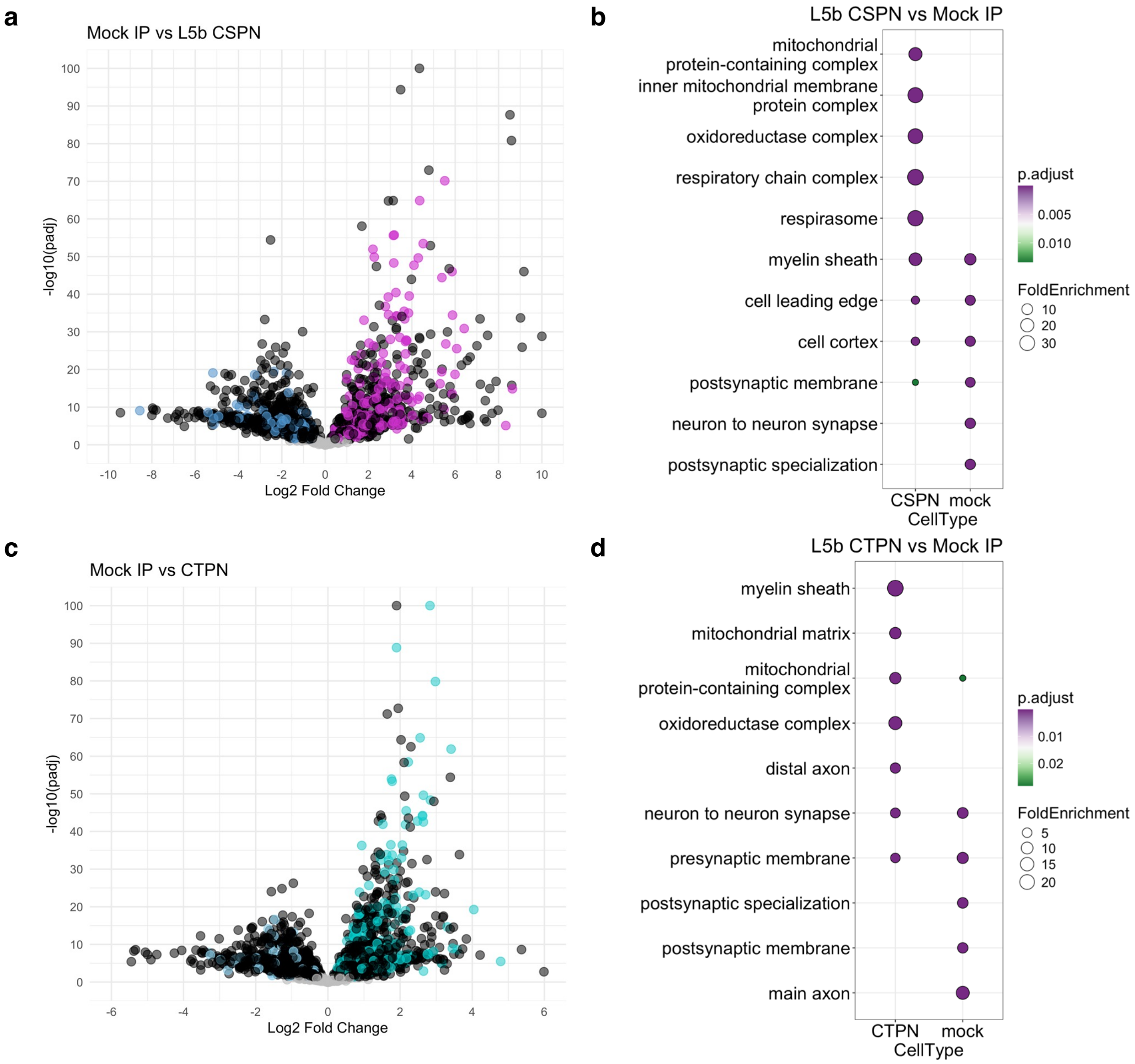


**Supplementary Figure 4. Enrichment of mitochondrial proteins in TOM-Tag IPs relative to mock controls. a**, Volcano plot of differential protein abundance in CSPN TOM-Tag IPs versus mock IPs. Magenta points indicate MitoCarta3.0 proteins significantly enriched in CSPN IPs; light blue points are MitoCarta proteins enriched in mock IPs; black points represent significantly different proteins not annotated in MitoCarta. **b**, GO cellular component enrichment analysis of proteins enriched in CSPN IPs compared to mock IPs. **c**, Volcano plot of differential protein abundance in CTPN TOM-Tag IPs versus mock IPs. Cyan points represent MitoCarta3.0 proteins significantly enriched in CTPN IPs; light blue and black points are defined as in (**a**). **d**, GO cellular component enrichment analysis of proteins enriched in CTPN IPs compared to mock IP**.**

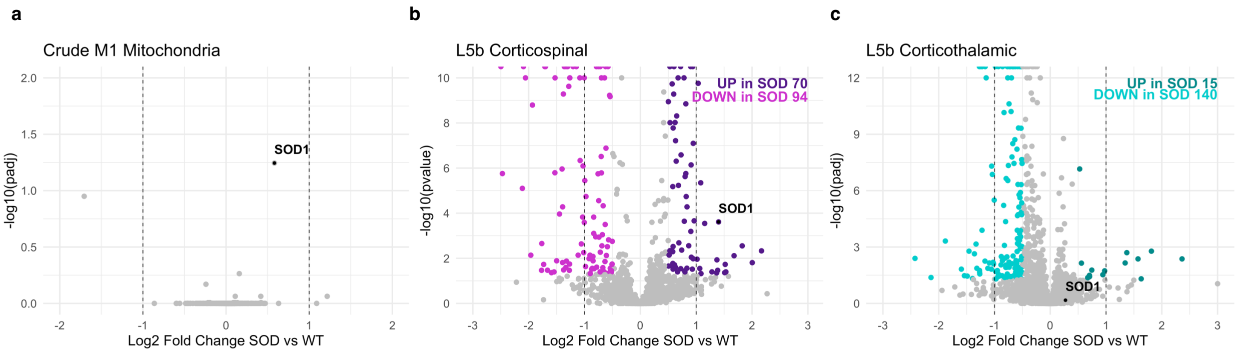


**Supplementary Figure 5. Differential mitochondrial proteomic responses to SOD1G93A in cortex and L5b projection neurons. a**, Volcano plot of differential protein expression in crude cortical mitochondria (SOD vs. WT). No proteins reached significance ($q < 0.05$) although SOD1 approached the threshold (q value= 0.057) and is labeled in black. **b**, Volcano plot of CSPN mitochondrial proteomes comparing SOD and WT samples. Significantly upregulated proteins are shown in purple and downregulated proteins in magenta. SOD1 is labeled in black. Counts of significant proteins from each group are displayed in the top right. **c**, Volcano plot of CTPN mitochondrial proteomes (SOD vs. WT). Significantly upregulated proteins are in dark cyan and downregulated in cyan with SOD1 indicated in black.
